# RNA-binding protein YBX1 promotes Type H vessels dependent bone formation in an m5C-dependent manner

**DOI:** 10.1101/2023.04.21.537774

**Authors:** Yu-Jue Li, Qi Guo, Wen-Feng Xiao, Ye Xiao

## Abstract

RNA Binding Proteins (RBPs) interact with RNA and ubiquitously regulating RNA transcripts during their life cycle. Previous works showed that RBPs play fundamental roles in the progression of angiogenesis-related diseases. However, the role of RBPs in skeletal endothelium-dependent bone formation and osteogenesis is unclear. Here, we show that RBP-Ybx1 was strongly reduced in bone vasculature from ovariectomy-induced osteoporotic mice. Endothelial cell-specific deletion of Ybx1 impaired CD31^hi^EMCN^hi^ endothelium morphology and osteogenesis and resulted in low bone mass, while its overexpression promoted angiogenesis-dependent osteogenesis and ameliorated bone loss in OVX mice. Mechanistically, Ybx1 deletion disrupted CD31, EMCN and BMP4 stability in an m5C-dependent manner and blocked endothelial-derived BMP4 release, thereby inhibiting osteogenic differentiation of BMSCs. Administration of recombinant BMP4 protein promoted osteogenic differentiation of BMSCs and restored impaired bone formation in Ybx1^iΔEC^ mice. Finally, tail vein injection of CD31-modified PEG-PLGA carrying sciadopitysin, a natural Ybx1 agonist, pharmacologically partially reversed CD31^hi^EMCN^hi^ vessels decline and improved the restoration of bone mass both in OVX and aging animals. These findings demonstrated the role of RBP-Ybx1 in angiogenesis-dependent bone formation and provided a novel therapeutic approach for ameliorating aging-related and postmenopausal osteoporosis.

## Introduction

Skeletal tissues are rich in blood vessels that consist of type H vessels (CD31^hi^EMCN^hi^) distributing in the metaphysis region with bud, arche, and column-shaped, and the highly branched type L vessels (CD31^lo^Emcn^lo^) distributing in the diaphysis region(Kusumbe *et al*, 2014). The skeletal blood vessels, specifically the type H vessels subtype, control bone homeostasis, repair, and pathobiological processes, and the disruption of the vascular network is associated with the progression of bone diseases including cancer and osteoporosis(Tuckermann & Adams, 2021). Osteoporosis is a systemic bone disease characterized by decreased bone mass, impaired bone microstructure, and fracture prone(Rachner *et al*, 2011). And postmenopausal osteoporosis is especially concerning because it significantly increases the fracture risk of older women. Recent research has found that the enrichment of skeletal type H vessels significantly declines in aging and ovariectomy (OVX)-induced mice and osteoporotic patients(Wang *et al*, 2017; Xie *et al*, 2014). On the one hand, the declined type H vessels contribute to reduced nutrient delivery, tissue metabolism, and the influx of calcium and phosphate; on the other hand, reduce the expression of vasogenic secreted proteins and block the crosstalk between endothelial cells and other cell types including pre-osteoprogenitors, osteoprogenitors, osteoblasts, chondrocytes, and hematopoietic stem cells and then impair bone homeostasis(McCarthy, 2006; Rafii *et al*, 2016; Tomlinson & Silva, 2013; Tuckermann & Adams, 2021). In the skeletal system, at the vessel buds’ proximal end, the bone mesenchymal stromal cells (BMSCs) are also surrounded by the relatively straight, column-shaped type H capillaries(Baccin *et al*, 2020). In addition, perivascular BMSCs interact with type H capillaries by secreting glioma-associated oncogene homolog 1 (GLI1) during bone development and defect healing(Chen *et al*, 2020). These suggested that BMSCs-derived instructive signals regulate endodermal cells. However, it is currently unclear whether and how skeletal endothelial cell networks are coupled with the differentiation of BMSCs.

RNA-binding proteins (RBPs) are a large class of proteins with RNA-binding properties and chaperone RNA throughout their lifetime and regulate RNA stability, modifications, and localization(Cook *et al*, 2011; Gerstberger *et al*, 2014; Xu *et al*, 2017). Several key RBPs have been reported to modulate the physiological and pathological angiogenesis by regulating the metabolism of messenger(m)RNAs encoding angiogenic modulators(Chang & Hla, 2011; Smith & Costa, 2022). However, the role of RBPs in type H vessel formation is not unclear. The Y-box-binding protein 1 (Ybx1), one of the RBP families, is involved in RNA splicing, stability, and translation control during the RNA lifecycle(Mordovkina *et al*, 2020). Evidence from other studies indicates YBX1 is implicated in physiological and pathological angiogenesis by regulating the translation and stability of VEGF-A and HIF1-α(Coles *et al*, 2004; Coles *et al*, 2002; Coles *et al*, 2005; El-Naggar *et al*, 2015; Smith & Costa, 2022). Due to the high expression of Ybx1 in tumor-associated vasculature, Ybx1 has also been studied to suppress tumor angiogenesis as a therapeutic target(Takahashi *et al*, 2010). Previously, we demonstrated that Ybx1 was regarded as a splicing factor to regulate BMSCs fat during aging(Xiao *et al*, 2023). However, the RBP-Ybx1 whether it participates in the regulation of angiogenesis of skeletal vessels, and the Ybx1 loss in endothelial cell lineage(s) is linked to osteoporotic disease.

In the present study, we found that endothelial-specific RBP-Ybx1 knockout mice showed decreased CD31^hi^EMCN^hi^ vessel density and osteogenic differentiation of BMSCs by disrupting the stability of CD31 and pro-osteogenesis factor BMP4. Administration of recombinant BMP4 restored the phenotype defects in Ybx1-deleted mice. Moreover, we constructed novel targeting-vessel nanoparticles, carrying sciadopitysin, which increase the CD31^hi^EMCN^hi^ vessels density and the secretion of BMP4 and promote bone formation in OVX and aged mice. Taken together, our study identified a potential therapeutic target and a novel approach to treat postmenopausal and aging-related osteoporosis.

## Methods

### Mice

All mice we used were on a pure C57/B6 background. The Cdh5-Cre transgenic mice were purchased from Jackson Laboratory; loxP-flanked Ybx1 mice were purchased from Cyagen Biosciences. Ybx1^+/-^ mice were crossed with Ybx1^+/-^ mice to generate Ybx1^+/+^ (wild-type, WT) and Ybx1^-/-^ (Ybx1^iΔEC^) mice. WT littermates of the same sex were used as controls. For endothelium-specific Ybx1 knockout experiments, eight female mice of 3 weeks were used for each group. C57BL/6J mice aged 20 months were bought from Charles River Animal Company (China). For the Ovariectomy (OVX) model, 4-week-old female mice were bilaterally ovariectomized or sham-operated. A month after Surgery, the OVX mice were treated with either sciadopitysin or rAAV2/9-Tie1-Ybx1. For sciadopitysin treatment, OVX mice were randomly divided into two groups and injected with sciadopitysin via the tail vein at a dose of 25 mg/kg every other day for 1 month. The sham group and control OVX group instead received vehicle reagents. For AAV treatment, OVX or sham mice were injected with 100μl rAAV2/9 (1 × 10^12^ cfu/ml) expressing GFP or Ybx1 via the tail vein for a month. Three days after the last injection, all mice were euthanized, and femurs and tibias were collected for follow-up experiments. All the mice were bred under specific-pathogen-free conditions at Laboratory Animal Research Center at Central South University and approved by the Medical Ethics Committee of Xiangya Hospital of Central South.

### Endothelial cell culture and functional assays

The human umbilical vein endothelial cell line (HUVEC) was bought from American Type Cell Collection (Manassas, VA, USA). The HUVEC was cultured in Endothelial Cell Medium (ScienCell, 1001) and was maintained at 37°C and 5% CO_2_ in a humidified atmosphere. ECM consists of 500 ml of basal medium, 25 ml of fetal bovine serum (FBS), 5 ml of endothelial cell growth supplement (ECGS) and 5 ml of penicillin/ streptomycin solution (P/S). For the cell viability assay, HUVECs were inoculated in a 96-well plate (1 × 10^4^ cells in 200 μl of culture medium per well) and cocultured, respectively, with natural small molecular compounds theaflavin 3-gallate (TF2A), Eriocitrin, Sciadopitysin, Isoginkgetin, and Bilobetin were purchasing from Target Molecule Corp (Target Mol) in an incubator containing 5% CO_2_ at 37 °C for 48 h. After that, the CCK8 reagents were added to the cell medium (10 μl for each well) and incubated for another 4 h. The absorbance was measured at 450 nm using an ultra micro-pore plate spectrophotometer (Epoch, Bio Tek). The OD values were calculated from three independent experiments. Endothelial cell migration assays were performed in 24-well plates. The treated or untreated HUVECs (1 × 10^4^) were seeded into the upper chamber with 200 μl medium and 600 μl serum-free medium was added to the lower chamber. At 16 h after seeding, the chambers were fixed and stained with 0.2% crystal violet (Beyotime Biotechnology, C0121). The non-migrating cells in the upper chamber were carefully removed using cotton swabs, the migrating cells were counted with Inverted Microscope Camera System (Leica) in five random fields of each filter. Endothelial cell tube-formation assay was conducted in 48-well plates pre-coated with BD Matrigel (CORNING, 354234). The treated or untreated HUVECs (2 × 10^4^) were seeded into wells and incubated for 12 h. The tube-like structures were imaged by Inverted Microscope Camera System (Leica) and calculated automatically using the ImageJ software.

### BMP4 treatment

BMP4 recombinant protein was produced with the help of Forevertech Biotechnologies Co.,ltd. The cultured primary BMSCs were treated with different concentrations of BMP4 and induced osteogenic or adipogenic differentiation according to the manufacturer’s instructions(Li *et al*, 2021). The BMSCs were treated with BMP4 with different concentrations for the duration of 24 h and subsequently lysed in trizol reagent for RT-qPCR analysis. The control and Ybx1-deleted mice were treated with BMP4 recombinant protein (3 μg/g) for 28 days. Three days after the last injection, all mice were euthanized, and femurs and tibias were collected for bone analysis as described earlier(Li *et al*, 2018).

### ELISA analysis

The HUVECs were infected with shControl or shYbx1 Adeno virus for 48 h. The cell mediums were collected and used to detect the TNFα levels by the TNFα ELISA kit (R&D Systems, MTA00B). All ELISA assays were performed according to the manufacturer’s instructions.

### Flow cytometry and cell sorting

The fresh femurs and tibias were dissected, crushed in cold Hanks′ Balanced Salt solution, and digested bone pieces with 1 mg/ml type IA collagenase (Sigma-Aldrich, SCR136) at 37°C for 20 min. After filtration and washing, the cells were counted and incubated for 45 min at 4°C with EMCN antibody (#2114518, Invitrogen, USA, 1:100) and CD31 antibody (#FAB6874G, R&D Systems, USA, 1:100), then washed and further incubated with DAPI (#ab285390, Abcam, UK, 1:2,000). Finally, the CD31^hi^EMCN^hi^ cells were acquired and demarcated on a BD FACScan cytometer (BD Immunocytometry Systems). The collected and sorted CD31^hi^EMCN^hi^ cells were lysed in trizol reagent for RT-qPCR analysis.

### Immunofluorescence and Immunocytochemistry of bone sections

For immunofluorescence staining, freshly dissected femora were fixed in 4% paraformaldehyde solution for 24 h, then decalcified in 0.5 M EDTA (pH 7.4) at 4°C for 10 days (4-month-old mice) or 21 days (21-month-old mice). Then the bones were incubated in cryoprotectant solution (10% sucrose and 1% polyvinylpyrrolidone) for 24 h at 4°C. After that, bone tissues were embedded in embedding solution (8% gelatin, 2% polyvinylpyrrolidone, and 20% sucrose) and transferred to a −80°C ultra-low-temperature freezer overnight. Then, the samples were then longitudinally oriented and cut to 15µm thickness before being stained with primary antibodies to mouse CD31 conjugated to Alexa Fluor 488 (R&D Systems, #FAB3628G, 1:100), CD31 (BD Pharmingen, #553370, 1:200), endomucin (#sc-65495, Santa Cruz Biotechnology, USA, 1:50), Ybx1 (#4202S, CST, USA, 1:200), VEGFA (#19003-1-AP, Proteintech, China, 1:200), Col 1 (#AB765P, Millipore, USA, 1:200), Osterix (#ab22552, abcam, 1:200), and Bmp4 (#30912, Forevertech Biotechnologies Co., Ltd, China, 1:500) overnight at 4°C. The next day, the samples were incubated with secondary antibodies conjugated with fluorescence tags at room temperature for 60 min. The fluorescent signals were captured via fluorescence microscopy. For immunocytochemistry staining, the bone tissues were embedded in paraffin and cut to 4 µm. The paraffin sections were de-waxed and stained with primary antibody OCN (#M173, Takara Bio, Japan, 1:100), and counterstained with Harris Hematoxylin.

### Quantitative real-time PCR analysis

Total RNA from cells was extracted using AG RNAex Pro Reagent (AG21102, Accurate Biology, China) and reverse transcribed into cDNA by using the 5× Evo M-MLV RT Premix (AG11706, Accurate Biology, China). Amplification reactions were set up in 20-µl reaction volumes containing 2× Pro Taq HS Probe Premix (AG11704, Accurate Biology, China) and the relative mRNAs expression level were normalized to the endogenous Gapdh expressions. Primer sequences are listed in Table S1.

### RNA sequencing and alternative splicing analysis

The total RNA was extracted and used to construct shYbx1 RNA-sequencing libraries as described previously(Xu *et al*, 2018). The cleanReads were aligned to GRCm38.p6, to obtain the location information on the reference genome and the specific sequence characteristics for the sequenced sample by hisat2. The htseq-count software was used to obtain the number of reads in each sample compared to the protein-coding gene. Then, FPKM algorithms were used to calculate gene expression, and DESeq2 software(Love *et al*, 2014) (BaseMean value was used to estimate the expression) was used to test Differentially expressed genes (DEGs) between shYbx1 groups and control groups with q value < 0.05 and log2 fold change > 0.58. GO enrichment aims to show the biological processes affected among different groups. Heatmaps were generated to show gene expression differences in different groups. For alternative splicing analysis (shYbx1) was performed by rMATS(Georgilis *et al*, 2018), splicing changes with a false discovery rate (FDR) < 0.05 and FDR > 0.1 were considered statistically significant.

### RNA-stability assay

The HUVECs infected with shYbx1 or shControl adenoviruses. After adenoviruses infection (24 h), the cells were treated with 5 μg/ml actinomycin D (S8964, Selleck Chemicals) or DMSO and collected at the indicated time points. The cells were lyzed in trizol reagent for RT-qPCR analysis.

### Western blot analysis

The HUVECs infected with shYbx1 or shControl adenoviruses or treated with sciadopitysin were lysed in RIPA buffer with Protease inhibitor for 30 minutes on ice. The lysates were centrifugated and transferred the cell supernatant to new tubes. Then, the cell supernatant was boiled with 6×SDS loading buffer, and separated by sodium dodecyl sulfatepolyacrylamide gel electrophoresis. Proteins were transferred to polyvinylidene difluoride membranes (IPFL00010, Millipore, USA) and incubated with specific antibodies. Primary antibodies were listed below: CD31 (66065-2-Ig, Proteintech, China, 1:1000), EMCN (sc-65495, Santa Cruz Biotechnology, USA, 1:500), YBX1 (4202S, CST, USA, 1:1000), BMP4 (P00205, Forevertech Biotechnologies Co.,ltd, China, 1:1000), Tubulin (#11224-1-AP, Proteintech, China, 1:1000). Secondary anti-mouse/rat/rabbit HRP-conjugated antibodies were subsequently applied.

### Cross-Linking and Immunoprecipitation High Throughput Sequencing

The cross-linking and immunoprecipitation experiments were performed with a CLIP Kit (BersinBio, Bes3014-2) according to the manufacturer’s instructions. Briefly, HUVECs were treated with 4-Thiouridine with a final concentration of 100 μM for 16 h and irradiated with 365 nm UV-C light (0.15 J/cm^2^) to covalently cross-link proteins / nucleic acids in vitro. The cross-linked cells were lysed in cell lysis buffer with DTT and Protease inhibitor for 10 minutes on ice. After centrifuging the cells at 13000 g for 5 minutes at 4°C, the supernatant was collected and incubated overnight at 4°C with ~5 µg YBX1 or IgG antibody and magnetic beads. To re-suspend the immunoprecipitates in IP wash buffer containing RNase T1 and incubate in a 22°C water bath for 15 min after washing the magnetic beads with IP wash buffer. Then, the immunoprecipitates were incubated with DNase I at 37°C for 15min. After that, immunoprecipitates were digested by Proteinase K and the bound RNA was extracted with phenol:chloroform:isoamyl alcohol according to the manufacturer’s instructions and then subjected to PCR amplification. The PCR products were used to construct libraries and the libraries were sequenced on the Illumina NovaSeq 6000 system by Wuhan Igenebook Biotechnology Co.,Ltd (Wuhan, China). Trimmomatic (version 0.38) filtered out low-quality reads from raw reads to generate clean reads. Bowtie2 (version:2.2.9) was used to map the clean reads to the GRCh38.p14 genome, and samtools (version 1.3.1) was used to remove potential PCR duplicates. CLIP-seq peaks were identified by Piranha (version 1.2.1) with the following parameters: “-p 0.00001 -z 100 -s -o”. Homer (version 4.9.1) was used for analysis of YBX1 binding motifs.

### m5C RIP and RNA-pull down

The m5C RIP was performed with an m5C RIP Kit from CloudSeq Biotech Inc. (GS-ET-003, China). Briefly, a total of ~100 µg RNA was digested in 1×Fragmentation Buffer, incubated at 70°C for 6min, and stopped the reaction by adding Stop Buffer. The RNA fragments were precipitated and re-dissolved in RNA-free water and incubated with PGM magnetic beads coupled with m5C antibody for 60min at 37°C. After that, the immunoprecipitates were washed with RLT Buffer and anhydrous ethanol three times. Finally, the binding RNA was eluted with RNA-free water and reverse transcribed into cDNA with random primers (AG11706, Accurate Biology, China). Semi-quantitative PCR was performed with BMP4, CD31, and an EMCN special primer. For the RNA-pull down experiment, BMP4 and CD31 specific biotinylated probes were synthesized by Sangon Biotech (Shanghai) Co., Ltd. and incubated with Ybx1 recombinant protein (NBP2-30101, Novus) at 4°C for 3 hours. The protein/nucleic acid complex was combined with streptomycin magnetic beads and pulled down on a magnetic stand. The beads were washed with 1× PBS four times and then subjected to sodium dodecyl sulfatepoly acrylamide gel electrophoresis after boiling with 1× SDS loading buffer.

### µCT analysis

Femuro were dissected from all the experimental groups of mice. After removing the attached soft tissue thoroughly, the femurs were fixed in 4% paraformaldehyde for 24 h. The fixed samples were scanned using high-resolution μCT (Skyscan 1172, Bruker microCT, German). The basic parameters of the scanner included x-ray energy tube voltage of 65 kV, a current of 153 μA, and a resolution of 15 µm per pixel. After scan, the image reconstruction software (NRecon, version 1.6, Bioz, USA), data analysis software (CT Analyser, version 1.9, Bruker microCT, German) and three-dimensional model visualization software (μCT Volume, version 2.0, Bruker microCT, German) were used to analyze the parameters of the distal femoral metaphyseal trabecular bone. And a series of planar cross-section images were generated in this process. 5% of femoral length below the growth plate was selected for microarchitecture analysis. The following parameters were calculated to describe the subchondral trabecular bone microarchitecture including trabecular bone volume per tissue volume, trabecular number, trabecular separation, and trabecular thickness.

### PEG-PLGA nanoparticles

10 mg of sciadopitysin was dissolved in 10 mL of acetone:chloroform (7:3) solution and 1 ml was added to a 250 mL round bottom flask. Add 130 μl of 100 mg/mL of PEG-PLGA (Shanghai Ponsure Biotech, Inc) chloroform solution and 200μl of 10 mg/mL of NH2-PEG-PLGA (Shanghai Ponsure Biotech, Inc) chloroform solution and mix well. Then add another 3 mL of chloroform solution and mix well. Evaporate the above liquid to form a thin film under reduced pressure using a rotary evaporator (N-1300D-WB, EYELA, Japan). The speed of rotation is 180 rpm/min and the pressure is reduced from 150 mbar to 100 mbar, 60 mbar, 25 mbar. Add 2 ml of ultrapure water to the thin film and hydrate on the rotary evaporator under a water bath at 35 °C for 15 min. The speed is 180 rpm/min and the pressure is 500 mbar. Use ultrasonic processor (JY98-IIIDN, scientz, China) to sonicate the above products. The ultrasonic power was 45.5 kW and the duration of ultrasonication was 3 min, with a 2 s pause for every 10 s of ultrasonication. After sonication, the free unencapsulated drug was removed by centrifugation at 3000 g for 5 min, and the supernatant was the nanoparticles.

### TRAP staining

For osteoclast differentiation analyses in vitro, we isolated bone marrow monocytes (BMMs) from mouse femur bones. BMMs were seeded into 24-well plates at a concentration of 1.5 × 10^4^ cells per well. Cells were stimulated with 100 ng/ml RANKL (462-TEC-010, Novus Biologicals, USA) and 50 ng/ml M-CSF (HZ-1192, Proteintech, China) combining with Ybx1^flox/flox^, Ybx1^iΔEC^ endothelial cell conditioned medium, TNFα neutralizing antibody (50349-RN023, Sino biological inc. China) for 6 day, as previously described(Peng *et al*, 2022). Osteoclasts were fixed and stained using the TRAP staining kit (294-67001, FUJIFILM Wako Pure Chemical Corporation, Japan). The differentiation experiments were conducted in triplicate.

### Statistical analyses

Statistical analyses were performed with SPSS 20.0. The data are presented as mean ± SEM. Two-tailed Student’s t test or one-way analysis of variance (ANOVA) tests with Tukey’s multiple comparison tests is used to assess statistical significance. P<0.05 was used to define statistical significance and indicated by ‘*’; P < 0.05 were indicated by ‘**’; P < 0.01 were indicated by ‘***’; P < 0.001 were indicated by ‘****’. The experiments were randomized and repeatable.

## Results

### 1. RNA binding protein Ybx1 is significantly reduced in CD31^hi^EMCN^hi^ endothelial cells of ovariectomy mice

To investigate Ybx1 expression in bone endothelium, we performed ovariectomy surgery (OVX) to investigate events under pathological conditions. Micro-computed tomography (micro-CT) and histomorphometric analysis showed a significant decrease in bone mass (bone volume over total volume; BV/TV) and thickness of trabeculae in OVX mice relative to sham controls (12-week-old) (Fig 1 A and B). And flow cytometric quantification showed a strong reduction in CD31^hi^Emcn^hi^ endothelial cells in OVX mice compared to the sham group (Fig 1 C-E). Meanwhile, Ybx1 mRNA was decreased in CD31^hi^Emcn^hi^ endothelial cells (ECs) fluorescence-activated cell (FACS)-sorting from flushed sham group femur in comparison to extracted OVX samples by real-time quantitative PCR (RT-qPCR) analysis (Fig 1 F). Likewise, immunofluorescence of femur sections with specific antibodies displayed weak CD31 and Emcn (Fig 1 G-I) staining and slightly lower Ybx1 expression (Fig 1 J). These observations suggested that Ybx1 protein is dynamically controlled during aging. Some earlier studies highlighted the potential role of Ybx1 in angiogenic blood vessel growth(El-Naggar *et al*., 2015; Xue *et al*, 2020). Endothelial migration and capillary tube formation were found to be impaired in Ybx1 knockdown groups when compared to controls (Appendix Fig S1 A-D). To further identify potential pro-angiogenic factors regulated by Ybx1 in bone endothelium, we performed RNA-sequencing (RNA-seq) transcriptional profiling. Analysis of the RNA-seq data identified 873 mRNAs with decreased translation and 1746 mRNAs with up-regulation (Fig 1 K). Gene ontology (GO) analysis demonstrated that angiogenesis and osteogenesis-associated differentially expressed genes were enriched and comprised the top 15 GO categories in the Ybx1 knockdown group compared to the control (Fig 1 L and M). RT-qPCR data further validated that Ybx1 knockdown decreased the levels of pro-angiogenic genes while anti-angiogenic genes showed up-regulation (Appendix Fig S1 E). These findings suggested a key role of Ybx1 in angiogenesis and suggested it might be involved in the induction of bone endothelium.

**Fig 1.**
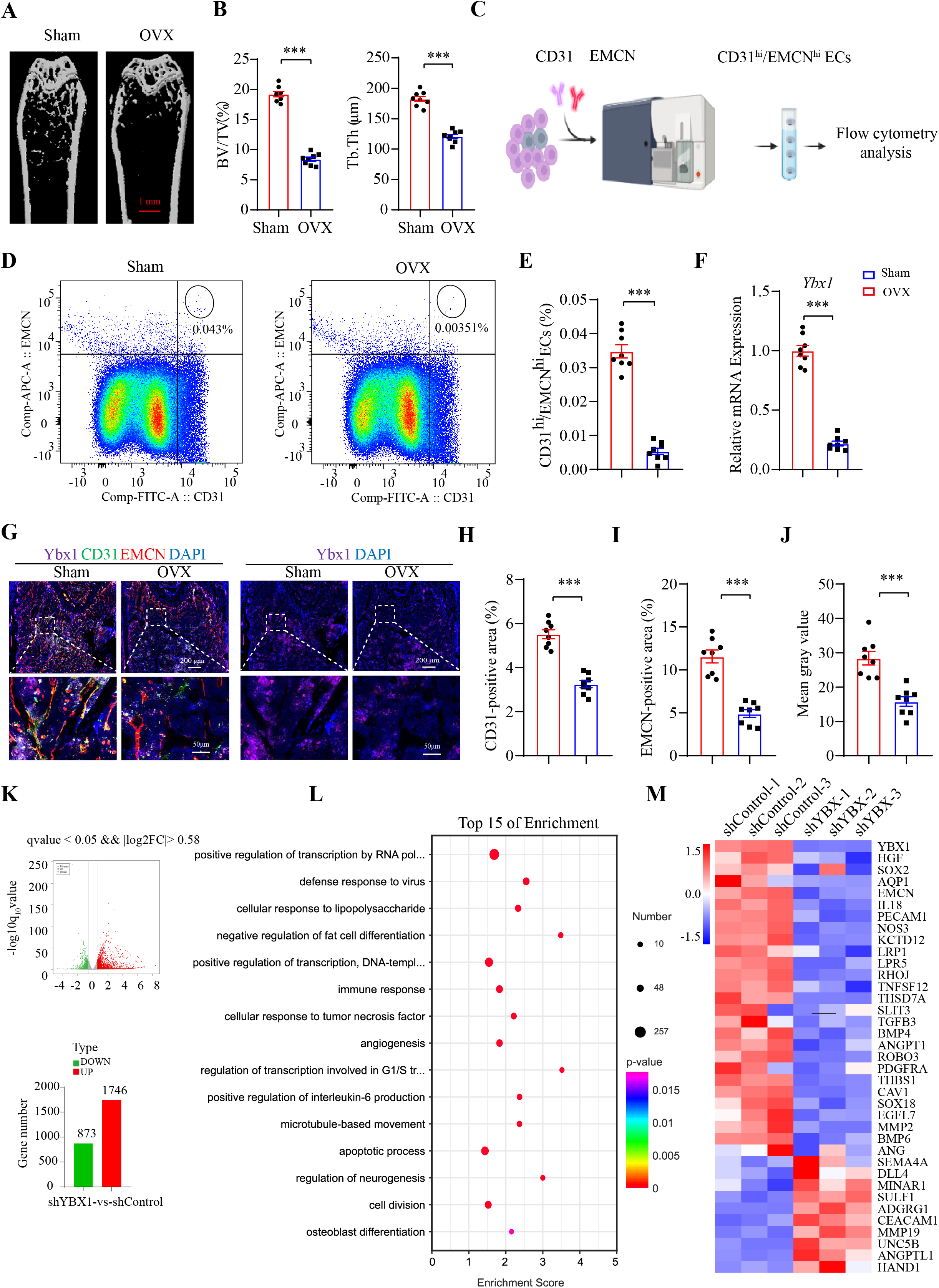
Ybx1 is reduced in CD31^hi^EMCN^hi^ endothelial cells of OVX mice. A-B. Representative µCT images (A) and quantitative µCT analysis (B) of femur of 4-month-old sham-operated and OVX mice. C-E. FACS analysis dot plot (C and D) and quantification (E) of CD31^hi^EMCN^hi^ ECs from sham-operated and OVX mice. F. RT-qPCR analysis of Ybx1 expression in CD31^hi^EMCN^hi^ ECs from sham-operated and OVX mice. G-J. Representative images (G) and quantification (H-J) of YBX1 (purple) in CD31 (green) -and EMCN (red) -stained femora from sham-operated and OVX mice. K-M. RNA-sequencing data from HUVECs with knockdown of YBX1. K. Among differentially expressed genes in HUVECs infected with shYBX1 and shControl adenovirus, 1746 genes were upregulated (red) and 873 were downregulated (green). L. Gene ontology (GO) analysis revealed enrichment of biological processes among the differentially expressed genes. M. Heatmap of differentially expressed genes related to angiogenesis. Data are shown as the mean ± SEM. ***, P < 0.001 by student’s t test.

### 2. Endothelial Ybx1 deletion impairs CD31^hi^EMCN^hi^ endothelium formation and bone formation

To further unveil the role of Ybx1 in bone endothelium, Ybx1 mutant mice (Ybx1^iΔEC^) with endothelial specific ablation of Ybx1 were generated by combining loxP-flanked Ybx1 alleles (Ybx1^lox/lox^) with Cdh5-Cre transgenics. RT-qPCR analysis of FACS sorted femur ECs showed a significantly knock-out efficiency of Ybx1 in Ybx1^iΔEC^ mice compared with control littermates (4-week-old) (Fig 2 A). The type H vessel (CD31^hi^EMCN^hi^) couples angiogenesis with osteogenesis and the decrease in richness of CD31^hi^EMCN^hi^ endothelium was closely integrated with aging-associated diseases such as postmenopausal osteoporosis(Huang *et al*, 2018; Kusumbe *et al*., 2014; Xie *et al*., 2014). To investigate the effect of Ybx1 on the richness of CD31^hi^EMCN^hi^ endothelium, type H endothelial cells were sorted and quantified by flow cytometry analysis. Our data showed that the amounts of type H endothelial cells sub-population were significantly decreased in the Ybx1^iΔEC^ metaphysis compared to littermate controls (Fig 2 B and C). And immunofluorescence analysis of Ybx1^iΔEC^ mutants revealed striking vascular defects. As showed in the experiment, the type H vessel density was strongly reduced and column/arch patterning as well as filopodia extension were disrupted in Ybx1^iΔEC^ mutant metaphysis (Fig 2 D and E). The phenotype was further supported by vascular endothelial growth factor A (VEGFA) immunostaining, a crucial regulator of both normal and pathological angiogenesis secreted by endothelial cells, perivascular cells, and mature/hypertrophic chondrocytes(Ferrara & Adamis, 2016; Fu *et al*, 2020) (Fig 2 F and G). This finding is consistent with previous report demonstrating that Ybx1 promotes angiogenesis by upregulation of VEGFA(Gao *et al*, 2021).

**Fig 2.**
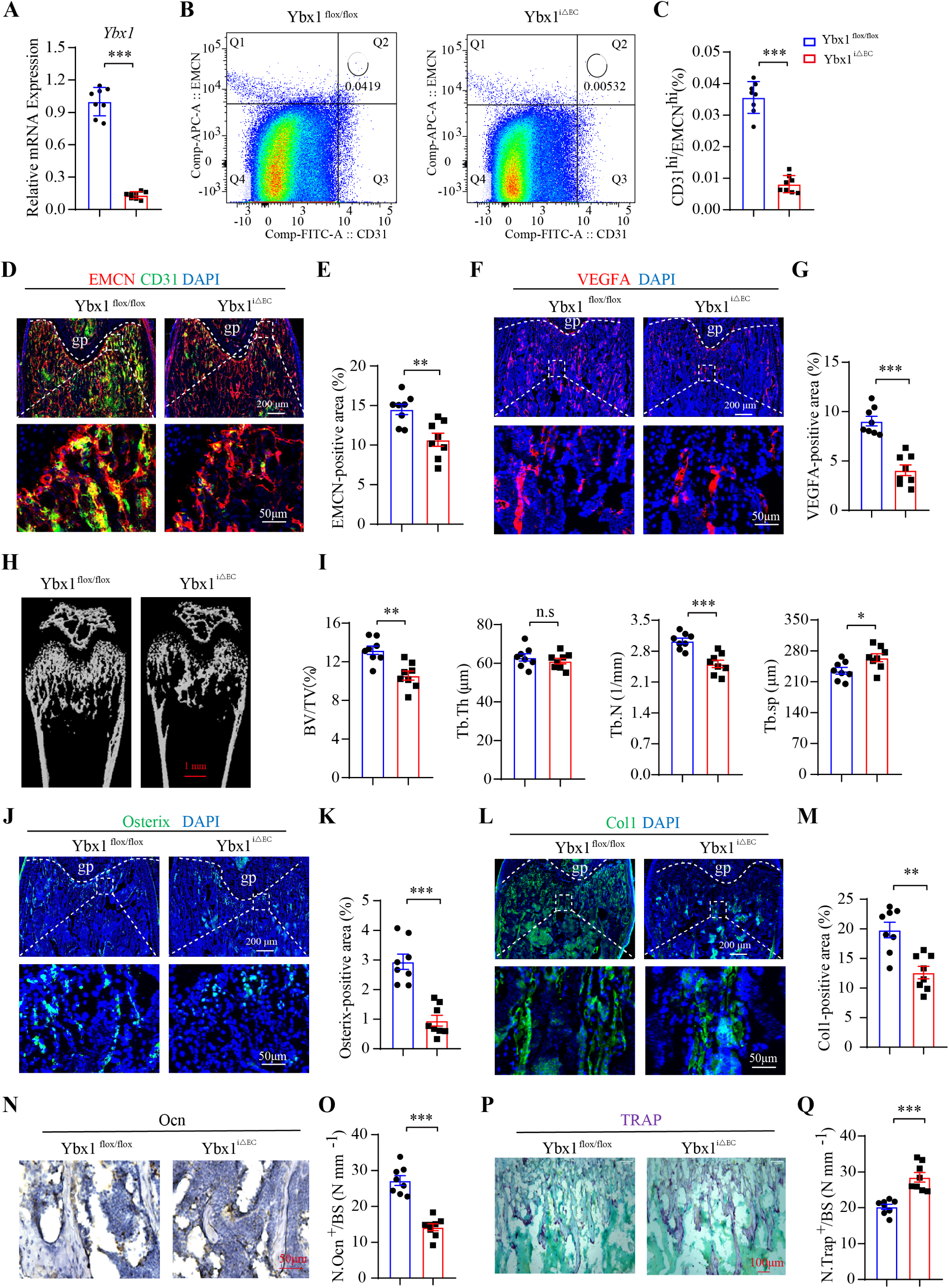
Endothelial Ybx1 deletion impairs CD31^hi^EMCN^hi^ endothelium formation and bone formation. A. RT-qPCR analysis of Ybx1 expression in CD31^hi^EMCN^hi^ ECs from endothelial-specific Ybx1 knockout mice (Ybx1^iΔEC^) and their littermate controls (Ybx1^flox/flox^). B-C. FACS analysis dot plot (B) and quantification (C) of CD31^hi^EMCN^hi^ ECs from Ybx1^iΔEC^ and Ybx1^flox/flox^ mice. D-E. Representative images (D) and quantitation (E) of CD31 (green) and EMCN (red) immunostaining in femora of juvenile 4-week-old Ybx1^iΔEC^ and Ybx1^flox/flox^ mice. Scale bar, 50 μm. F-G. Representative images (D) and quantitation (E) of VEGFA (red) immunostaining in femora of 4-week-old YBX1^iΔEC^ and YBX1^flox/flox^ mice. Scale bar, 50 μm. H-I. Representative µCT images (H) and quantitative µCT analysis (I) of trabecular bone microarchitecture of 4-week-old YBX1^iΔ EC^ and YBX1^flox/flox^ mice. Scale bar, 1 mm. J-K. Representative images (J) and quantitation (K) of Osterix^+^ (green) immunostaining in femora of juvenile 4-week-old Ybx1^iΔEC^ and Ybx1^flox/flox^ mice. Scale bar, 50 μm. L-M. Representative images (L) and quantitation (M) of COL1 (green) immunostaining in femora of juvenile 4-week-old Ybx1^iΔEC^ and Ybx1^flox/flox^ mice. Scale bar, 50 μm. N-O. Immunohistochemical staining (N) and quantification (O) of Ocn^+^ cells (brown) in trabecular bone surfaces of juvenile 4-week-old Ybx1^iΔEC^ and Ybx1^flox/flox^ mice. Scale bar, 50 μm. P-Q. Representative images and quantification of TRAP^+^ cells in trabecular bone surfaces of juvenile 4-week-old Ybx1^iΔEC^ and Ybx1^flox/flox^ mice. Scale bar, 100 μm. n = 8 mice in each group. Data are shown as the mean ± SEM. *, P < 0.05; **, P < 0.01; ***, P < 0.001 by student’s t test. Tb. BV/TV, trabecular bone volume per tissue volume; Tb. N, trabecular number; Tb. Sp, trabecular separation; Tb. Th, trabecular thickness.

Disrupted bone endothelium is accompanied with bone mass loss(Kusumbe *et al*., 2014). To investigate whether loss of Ybx1 influences bone mass in vivo. Micro-computed tomography (micro-CT) and histomorphometric analysis showed Ybx1^iΔEC^ mice with a severe osteoporosis phenotype, including a significant lower trabecular bone mass and density (that is, bone volume over total volume; BV/TV), trabecular thickness (Tb. Th.), trabecular number (Tb. N.) and expanded trabecular separation (Tb. Sp) (Fig 2 H and I). It has been previously reported that Osterix^+^ osteoprogenitors cells decreasing with age and have been identified to be responsible for bone loss and bone fracture(Smith *et al*, 1975). As expected, a remarkable decline in osteoprogenitors, co-coupling with bone vessels, was seen in 4-week-old Ybx1^iΔEC^ mice (Fig 2 J and K). Likewise, collagen type1α^+^ osteoblastic cells were reduced significantly (Fig 2 L and M). In addition, Osteocalcin (Ocn) immunostaining showed that Ybx1 mutants with fewer mature osteoblasts (Fig 2 N and O) whereas with appreciable changes in bone-resorbing osteoclasts compared to littermate controls (Fig 2 P and Q). Thus, these findings demonstrate that endothelial Ybx1 deletion elicits impairment of Type H endothelium formation and bone formation, whereas it activates bone resorption.

### 3. AAV2/9. sup-Tie1-Ybx1 treatment alleviates ovariectomy-induced bone loss

As we reported, the loss of Ybx1 in type H vessels is supported as a main reason for the defect of bone endothelium and bone formation. Importantly, in a gain-of-function assay, we showed that over-expression of Ybx1 promoted capillary tube formation and endothelial migration in a concentration-dependent manner (Appendix Fig S2 A-C). Next, we conducted an in vivo experiment to evaluate the effect of endothelial cell specific Ybx1 overexpression on OVX-induced bone loss. OVX mouse models were generated by cutting the ovaries from 4-week-old female mice and sham-operation (Sham) mice were only performed with surgery. Adeno-associated virus (AAV) 2/9 sup-Tie1-Ybx1 was injected into bone vessels by tail vein injection at 4 weeks post-surgery. Post-surgery at 8 weeks, the femurs were collected and then immunostained with specific GFP and EMCN antibodies. The result showed that a strong green fluorescence signal was detected in the bone endothelium (Appendix Fig S2 D). RT-qPCR analysis of femur ECs sorted from Ybx1 over-expressing mice (Ybx1^iOE-EC^) revealed a boosted Ybx1 mRNA level compared with control mice (Fig 3 A). And flow cytometric quantification analysis confirmed an OVX-induced reduction of CD31^hi^Emcn^hi^ endothelial cells, whereas endothelial cell numbers increased in Ybx1^iOE-EC^ mice (Fig 3 B and C). Likewise, immunostaining of femur sections showed Ybx1 overexpressing mutants (Ybx1^iOE-EC^) partly rescued OVX-induced decrease in the richness of CD31^hi^EMCN^hi^ endothelium and the defect of column/arch patterning and filopodia extension (Fig 3 D). As well, the VEGFA density was efficiently recovered in OVX mice treated with AAV 2/9 sup-Tie1-Ybx1 (Fig 3 E). Micro-CT analysis and histomorphometric analysis showed that AAV 2/9 sup-Tie1-Ybx1 injection markedly rescued bone loss in OVX mice (Fig 3 F and G). Likewise, the decrease of Osterix^+^ osteoprogenitors and osteoblast number in femur metaphysis of OVX mice were also partly improved following AAV 2/9 sup-Tie1-Ybx1 treatment (Fig 3 I and J) whereas the treatment had an opposite effect on osteoclast number (Fig 3 K). These data suggested that defective angiogenesis and osteogenesis and abnormal activation of osteoclastogenesis in OVX mice were partly reversed by treating with Ybx1 overexpression AAV 2/9 in endothelial cells.

**Fig 3.**
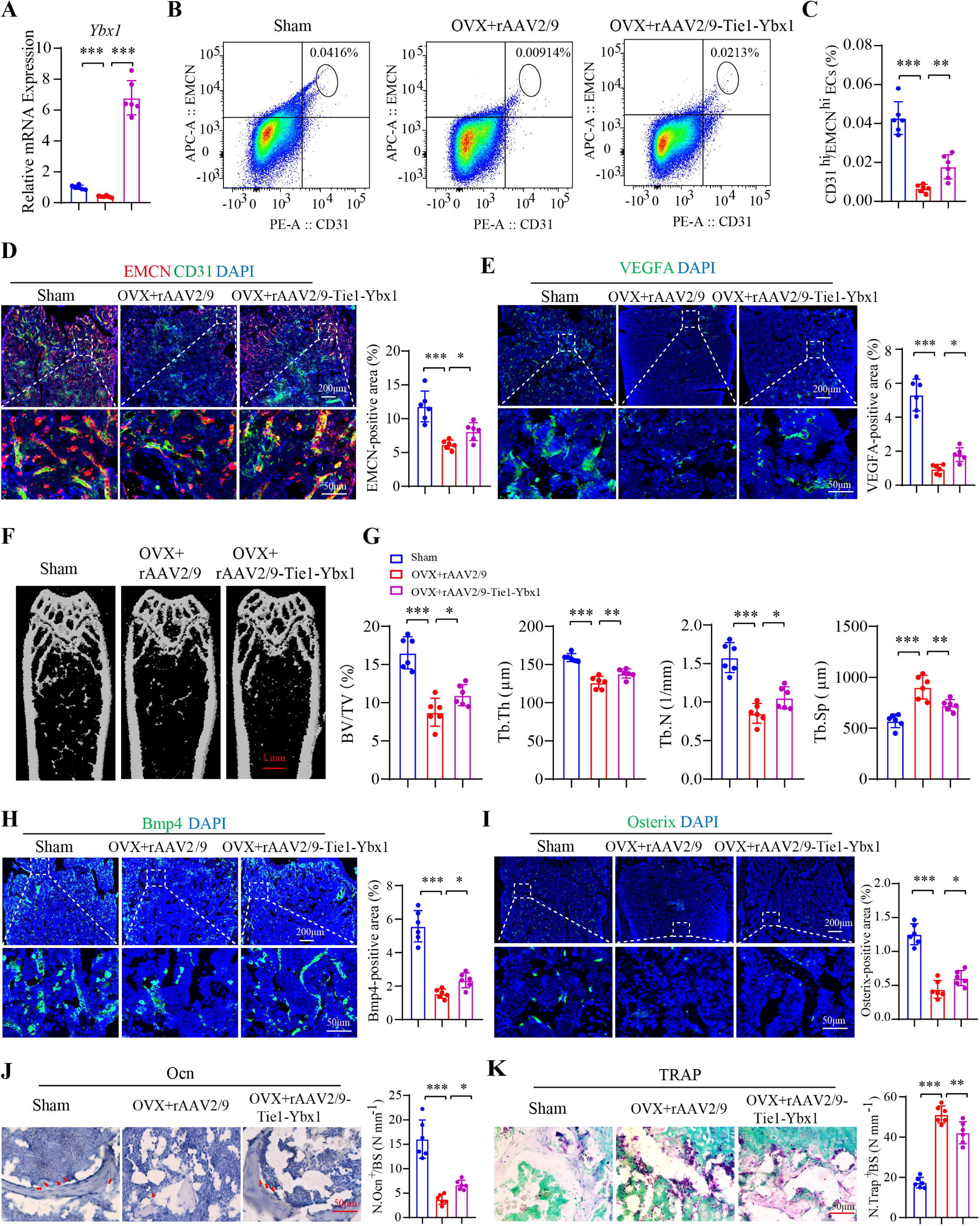
AAV2/9. sup-Tie1-Ybx1 treatment alleviates ovariectomy-induced bone loss. A. RT-qPCR analysis of Ybx1 expression in CD31^hi^EMCN^hi^ ECs from sham, OVX, and OVX mice injected with AAV2/9. sup-Tie1-Ybx1. B-C. FACS analysis dot plot (B) and quantification (C) of CD31^hi^EMCN^hi^ ECs from sham, OVX, and OVX mice injected with AAV2/9. sup-Tie1-Ybx1. D. Representative images and quantitation of CD31 (green) and EMCN (red) immunostaining in femora of sham, OVX, and OVX mice injected with AAV2/9. sup-Tie1-Ybx1. Scale bar, 50 μm. E. Representative images and quantitation of VEGFA (green) immunostaining in femora of sham, OVX, and OVX mice injected with AAV2/9. sup-Tie1-Ybx1. Scale bar, 50 μm. F-G. Representative µCT images (F) and quantitative µCT analysis (G) of trabecular bone microarchitecture of sham, OVX, and OVX mice injected with AAV2/9. sup-Tie1-Ybx1. Scale bar, 1 mm. H. Representative images and quantitation of Bmp4 (green) immunostaining in femora of sham, OVX, and OVX mice injected with AAV2/9. sup-Tie1-Ybx1. Scale bar, 50 μm. I. Representative images and quantitation of Osterix^+^ (green) immunostaining in femora of sham, OVX, and OVX mice injected with AAV2/9. sup-Tie1-Ybx1. Scale bar, 50 μm. J. Immunohistochemical staining and quantification of Ocn^+^ cells (brown) in trabecular bone surfaces of sham, OVX, and OVX mice injected with AAV2/9. sup-Tie1-Ybx1. Scale bar, 50 μm. K. Representative images of TRAP staining and Quantification of TRAP^+^ cells in trabecular bone surfaces of sham, OVX, and OVX mice injected with AAV2/9. sup-Tie1-Ybx1. Scale bar, 50 μm. n = 6 mice in each group. Data are shown as the mean ± SEM. *, P < 0.05; **, P < 0.01; ***, P < 0.001 by one-way ANOVA. Tb. BV/TV, trabecular bone volume per tissue volume; Tb. N, trabecular number; Tb. Sp, trabecular separation; Tb. Th, trabecular thickness.

### 4. YBX1 depletion leads to decreasing CD31 and EMCN stability in an m5C-dependent manner

To distinguish between direct and indirect YBX1 angiogenesis and osteogenesis-associated targets, and to investigate the detailed context-dependent rules for YBX1 regulation, we performed clip-seq (crosslinking-immunoprecipitation and high-throughput sequencing). We used bowtie2 (version:2.2.9) to map the sequenced reads to the known Gene set from NCBI GRCh38.p14 genome database and identified 819 significant peaks number using an FDR of <0.05 and found that most of the peaks are localized in exons (27.1%) and introns (18.97%) (Fig 4 A). Notably, a sizable fraction of peaks was mapped to the 3’-UTR region (14.9%) (Fig 4 A). Of the target genes assayed, we found that 64 common targets out of 4310 YBX1 knockdown altered targets and out of 284 YBX1 Clip-seq targets were bound by RNA-seq and Clip-seq data (Fig 4 B). We were surprised to find that the most significantly enriched target genes involve angiogenesis, including pro-angiogenesis genes - ANGPT2, CD31, EMCN, RUNX1, and THBS1 (Fig 4 B). It suggested that YBX1 affects the angiogenesis of endothelial cells by regulating pro-angiogenesis genes expression. It is worth noting that the majority of the peaks are clustered in a specific region, implying that multiple YBX1-binding events are involved in regulating pre-mRNA splicing. Because YBX1 has been reported to exhibit specific biological functions in the progression of pre-mRNA splicing and mRNA stabilization through direct RNA binding(Jayavelu *et al*, 2020; Yang *et al*, 2019). To test whether YBX1 regulates pre-mRNA splicing of candidate targets, we initially calculated the alternative splicing changes in RNA-seq data by measuring “false discovery rate” (FDR) and “IncLevelDifference” values. Exon skipping (ES) accounted for approximately 68% of all AS events, with the others including alternative 5’ splice site, alternative 3’ splice site, mutually exclusion exon, and retained intron in YBX1 knockout versus control with an FDR < 0.05 and IncLevelDifference > 0.01 (Appendix Fig S3 A and B). However, we noticed that no aberrant splicing was shown for EMCN and CD31 (Appendix Fig S3 C - E). It suggested that YBX1 affects angiogenesis and osteogenesis in an alternative splicing-independent way. Previous studies reported that Ybx1 regulates mRNA stability by binding to the 3’-UTR region(Yang *et al*., 2019). So, we next focused on 3’-UTR regional clustered YBX1-binding events by identifying peaks above the gene-specific. We first determined the known and novel overrepresented motifs in YBX1-binding clusters by using the Homer motif analysis software. Our data showed that about 24.78% of clusters contain the top-scored motif CCCCUC, and about 18.67% of clusters contain the motif ACCACC (Fig 4 C). Obviously, most CLIP clusters were enriched in CC-rich hexamers. It is consistent with the fact that Ybx1 preferentially recognizes m5C-modified mRNAs in the 3’-UTR region and plays essential roles in maternal mRNA stability(Chen *et al*, 2019; Yang *et al*., 2019). Indeed, CLIP-Seq analysis revealed that YBX1 had significantly higher peak distributions on the 3’-UTR region of CD31 and EMCN (Fig 4 D-F). To further assess whether YBX1 binds to the three targets 3’-UTR, we performed m5C RNA Binding Protein Immunoprecipitation (RIP) and found YBX1 binds to CD31, and EMCN 3’-UTR region upon m5C IP but not IgG immunocomplexes (Fig 4 G). The result was confirmed by using biotin-labeling RNA-pull down analysis with a gene-specific probe containing the BMP4 and CD31 or MUT 3′UTR with a mutated CC-rich motif (m5C site) (Fig 4 H and I). We next determined the specific role of the YBX1 binding m5C site in regulating 3′UTR activation. The CD31 WT 3′UTR and MUT 3′UTR were constructed into pGL-4 vector, respectively and co-transfected with pCMV-YBX1 into HEK-293T cells. As expected, YBX1 substantially activated the luciferase activity of WT 3′UTR but not MUT 3′UTR (Fig 4 J), providing further evidence that YBX1 regulates mRNA levels by binding to its 3′UTR region specifically with m5C modification. Proteins that bind 3’ UTRs are associated with regulating mRNA stability, translation, and localization(Ji *et al*, 2011). To examine whether a loss of YBX1 expression affects the CD31 and EMCN mRNA stability, HUVECs cells were infected with YBX1 shRNA and negative shRNA for 2 days followed by treatment with actinomycin D for various time points. Quantitative RT-PCR showed that CD31 and EMCN mRNA levels were degraded faster upon actinomycin D treatment and the relative CD31 mRNA half-life (9.5 h) was reduced by 91.5% (Fig 4 L) and EMCN mRNA half-life (10.9 h) was reduced by 60.5% (Fig 4 M) in shYBX1-expressing HUVEC cells compared to shControl group. Furthermore, YBX1 knockdown significantly reduced CD31 and EMCN protein expression (Fig 4 N). Therefore, we uncovered that deletion of YBX1 impaired angiogenesis by disrupting CD31 and EMCN mRNA stability in an m5C-dependent manner.

**Fig 4.**
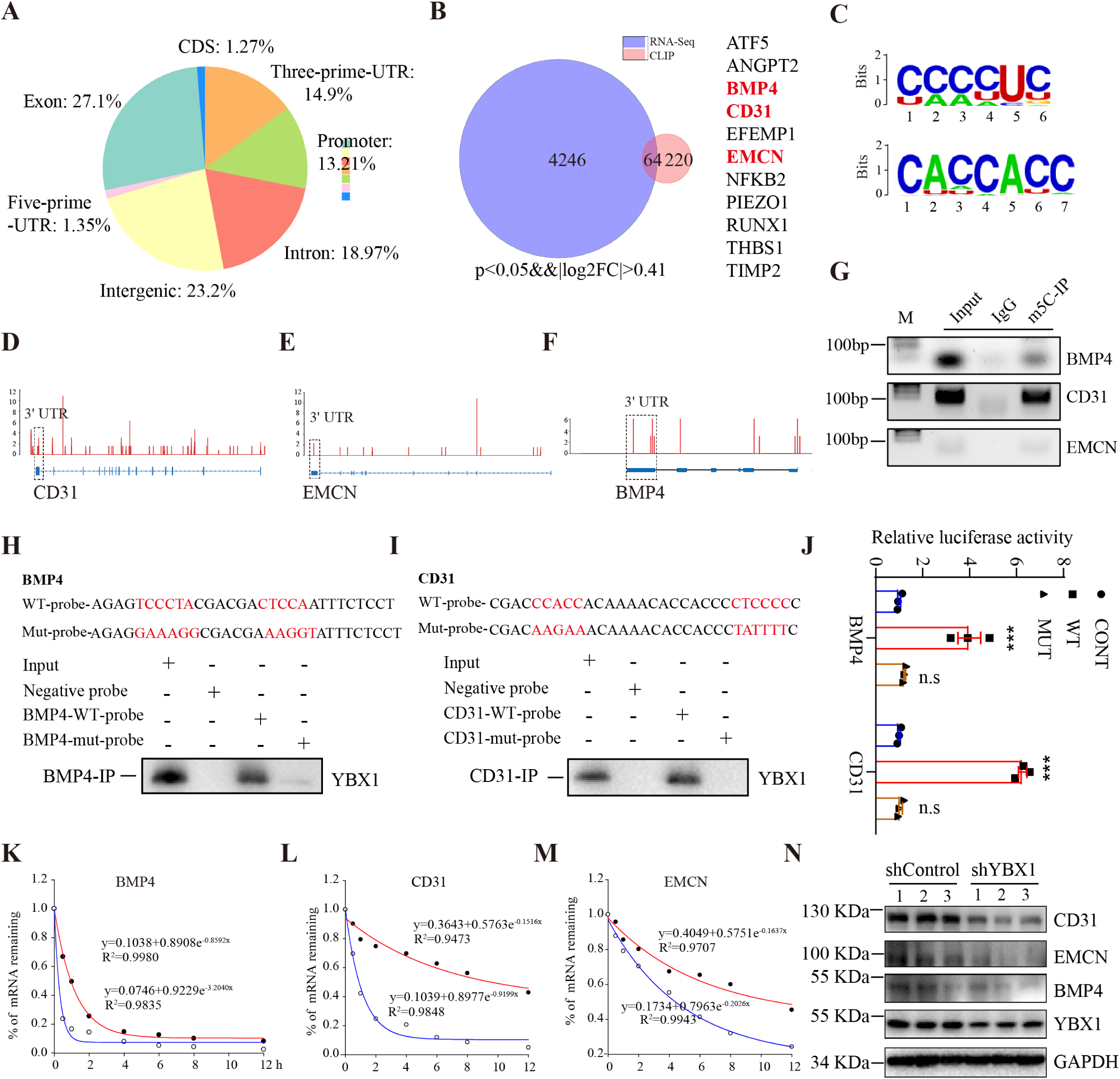
YBX1 depletion leads to decreasing CD31 and EMCN stability in an m5C-dependent manner. A. Genomic distribution of YBX1 CLIP-seq peaks. B. Venn diagram representing the overlap genes between YBX1 CLIP-seq targets and YBX1 knockdown RNA-seq targets. C. Top 2 ranked sequence motifs enriched in YBX1 CLIP-seq. D-F. Genomic view of YBX1 binding to CD31, EMCN and BMP4 loci. The frame area is 3′UTR. G. Semi-quantitative PCR showed RNA binding protein immunoprecipitates using m5C-RIP kit. H-I. RNA pull down analysis of binding between YBX1 protein and CD31 (or BMP4) - WT (MUT) - probe. J. Relative luciferase activity of HEK-293T cells transfected with different PGL-4 vectors and pCMV-YBX1. K-M. RT-qPCR analysis of the BMP4, CD31 and EMCN mRNA degradation rates. N. Western blotting analysis of the relative levels of CD31, BMP4, EMCN and YBX1 protein expression. n = 3 independent experiments. Data are shown as the mean ± SEM. ***, P < 0.001 by one-way ANOVA.

### 5. BMP4 secreting from bone vessel restores bone formation by coupling with BMSCs osteogenic differentiation

At the growth plate proximal end, Type H vessels are strongly associated with perivascular bone mesenchymal stromal cells (BMSCs) and osteoprogenitor cells in the metaphysis(Tikhonova *et al*, 2019). In addition, an impaired function of Type H bone vessels lost the promoting effect on BMSCs proliferation(Cui *et al*, 2022), which indicates Type H bone vessels may be involved in affecting BMSCs function. To investigate whether endothelial cell-specific deletion of YBX1 affects BMSCs function and understand how endothelial cell-specific deletion of YBX1 affects bone formation, we harvested conditioned medium from WT and YBX1^iΔEC^ BM-derived endothelial cells and co-cultured them with primary BMSCs. Conditioned medium from WT endothelial cells displayed an enhanced ability to induce osteogenic differentiation of BMSCs, whereas it inhibited adipogenic differentiation in comparison to the basic medium. In contrast, the effects were abolished in YBX1^iΔEC^ endothelial cells conditioned medium (Appendix Fig S4 A-C). Co-immunostaining of CD31 and leptin receptor (LepR) demonstrated that LepR-positive BMSCs were highly surrounded by bone endothelial cells and the amount of CD31 vessels, as well as the accompanying LepR-positive BMSCs, were dramatically reduced in the YBX1^iΔEC^ mice compared with their littermate control (Fig 5 A). Our results suggested that bone endothelial cells and BMSCs differentiation seem to be coupled. Endothelial cells control the behaviour of other cell types in the surrounding tissue by paraclinical molecular signals(Rafii *et al*., 2016). These results demonstrated that there is a soluble vessel-derived factor crosstalk between endothelial and BMSCs. To identify the potential factors regulated by YBX1 in endothelial cells, we screened the common targets derived from RNA-seq and Clip-seq data and identified the bone morphogenetic protein (BMP) family member BMP4, a pro-osteogenesis factor for coupling of angiogenesis and osteogenesis in bone development(Kusumbe *et al*., 2014; Langen *et al*, 2017). Because our previous results showed that YBX1 binds to the BMP4 3′UTR (Fig 4 D, G, and H) and regulates BMP4 mRNA stability in an m5C-dependent manner (Fig 4 J, K, and N). Immunofluorescence staining and quantification analysis showed that Ybx1^iΔEC^ mice with significantly lower BMP4 expression and weaker CD31 staining in comparison to littermate controls (Fig 5 B). In addition, we were surprised to find that BMP4 was expressed in the bone endothelium, strongly indicating that BMP4 is the potential soluble vessel-derived factor for coupling angiogenesis and BMSCs differentiation. To determine whether BMP4 involves in the regulation of BMSCs differentiation by bone endothelial cells in vitro, we treated primary BMSCs with BMP4 recombinant protein and induced osteogenic and adipogenic differentiation. RT-qPCR analysis showed that treatment of BMSCs with BMP4 led to upregulation of alkaline phosphatase (alp), ocn, and runx2 transcripts, and downregulation of fatty acid binding protein 4 (fabp4) and peroxisome proliferative activated receptor gamma (ppargγ) in a concentration-dependent manner (Fig 5 C and D). Meanwhile, BMSCs were treated with BMP4 to accumulate more calcium nodules and fewer fat drops by alizarin red and oil red O staining (Fig 5 E and F). It is consistent with its known role in bone formation that BMP4 promoted primary BMSCs differentiation into osteoblasts in vitro(Lee *et al*, 2011). We next assess whether administration of BMP4 restored the formation of trabeculae in endothelial cell-specific YBX1 deletion mice in vivo. BMP4 recombinant protein was daily injected into Ybx1^iΔEC^ mice (3 μg/g) for 4 weeks before analysis at day 28. In sharp contrast, administration of BMP4 recombinant protein rescued the structural alterations seen in the Ybx1^iΔEC^ metaphysis and bone vasculature (Fig 5 G and H), reduced bone marrow fat accumulation (Fig 5 I), enabled trabeculae formation (Fig 5 J), improved the normalized number of Lepr-positive BMSCs (Fig 5 K), but without detectable change in TRAP-positive osteoclasts (Fig 5 L). Our data demonstrates that bone vessel-derived factor BMP4 is actively involved in the coupling effect of endothelial YBX1 on bone angiogenesis and osteogenesis.

**Fig 5.**
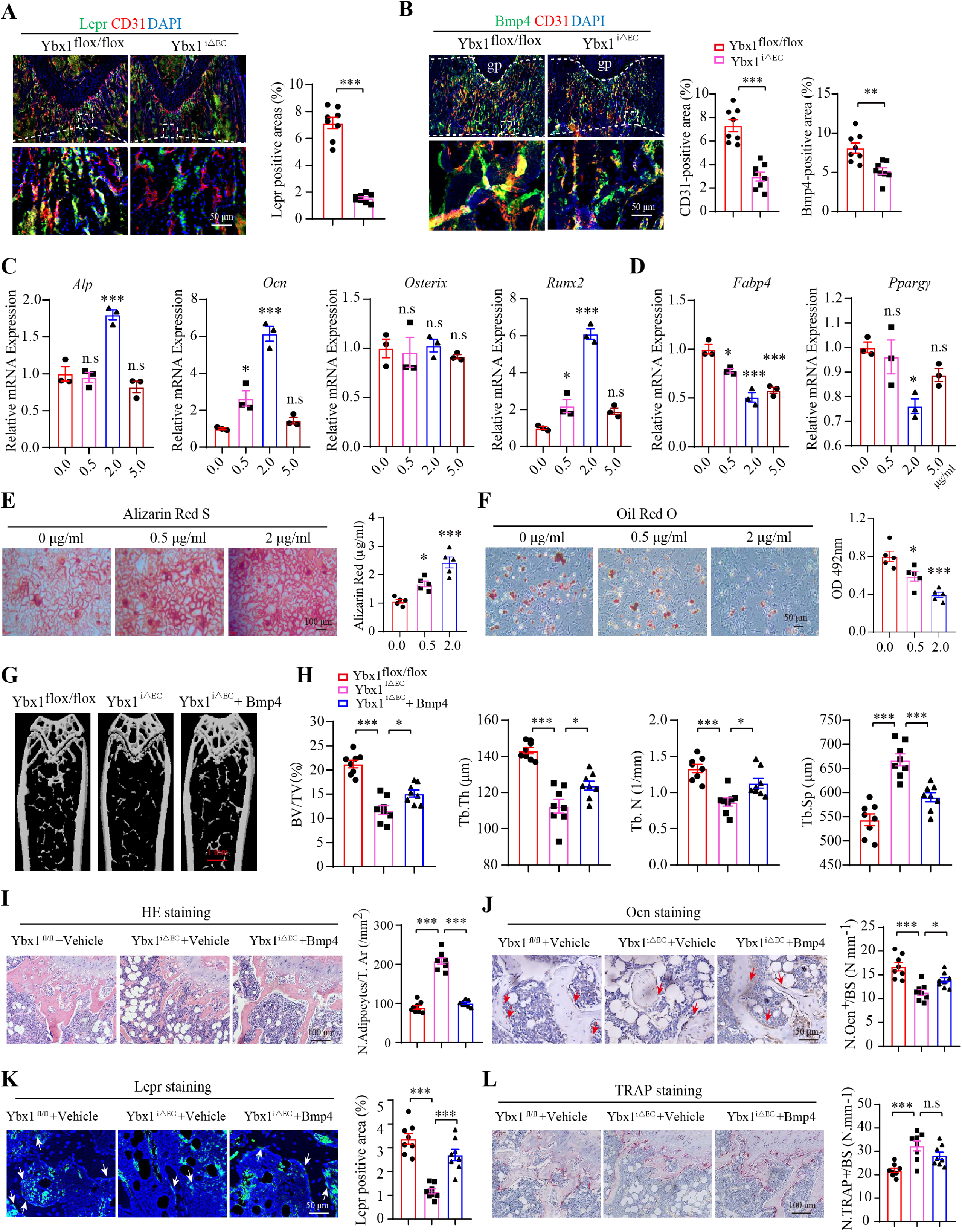
BMP4 recombinant protein restores bone formation by coupling with BMSCs osteogenic differentiation. A. Representative images and quantitation of Lepr (green) and CD31 (red) immunostaining in femora of juvenile 4-week-old Ybx1^iΔEC^ and Ybx1^flox/flox^ mice. Scale bar, 50 μm. B. Representative images and quantitation of Bmp4 (green) and CD31 (red) immunostaining in femora of juvenile 4-week-old Ybx1^iΔEC^ and Ybx1^flox/flox^ mice. Scale bar, 50 μm. C and D. RT-qPCR analysis of Alp, Ocn, Osterix, Runx2 expression levels (C) and Fabp4 and Ppargγ expression levels (D) in HUVECs treated with different concentrations of BMP4 recombinant protein. E. Representative images of Alizarin Red S staining and quantification of calcification of BMSCs treated with different concentrations of BMP4 recombinant protein. F. Representative images of Oil Red O staining and quantification of lipid formation of BMSCs treated with different concentrations of BMP4 recombinant protein. G-H. Representative µCT images (G) and quantitative µCT analysis (H) of trabecular bone microarchitecture of YBX1^flox/flox^, YBX1^iΔEC^ mice, and YBX1^iΔEC^ mice injected with BMP4 recombinant protein. Scale bar, 1mm. I. Representative images of H&E staining (Hematoxylin–Eosin Staining) in distal femora and quantification of the number of adipocytes related to the tissue area. Scale bar, 100 μm. J. Immunohistochemical staining and quantification of Ocn^+^ cells (brown) in trabecular bone surfaces of YBX1^flox/flox^, YBX1^iΔEC^ mice, and YBX1^iΔEC^ mice injected with BMP4 recombinant protein. Scale bar, 50 μm. K. Representative images and quantitation of Lepr (green) immunostaining in femora of YBX1^flox/flox^, YBX1^iΔEC^ mice, and YBX1^iΔEC^ mice injected with BMP4 recombinant protein. Scale bar, 50 μm. L. Representative images of TRAP staining and quantification of TRAP^+^ cells in trabecular bone surfaces of YBX1^flox/flox^, YBX1^iΔEC^ mice, and YBX1^iΔEC^ mice injected with BMP4 recombinant protein. Scale bar, 100 μm. n = 8 mice in each group. Data are shown as the mean ± SEM. *, P < 0.05; **, P < 0.01; ***, P < 0.001 by student’s t test (A and B) and one-way ANOVA (C-L). Tb. BV/TV, trabecular bone volume per tissue volume; Tb. N, trabecular number; Tb. Sp, trabecular separation; Tb. Th, trabecular thickness.

### 6. PEG-PLGA nanoparticles-carriers sciadopitysin acts as YBX1 agonists and enhances angiogenesis dependent bone formation in ovariectomy mice and aged mice

Evidence to date indicates that the enrichment of type H vessels is significantly lower in the bone of ovariectomy mice(Xie *et al*., 2014). Notably, our results demonstrated Ybx1 expression was slightly reduced, which correlates with the age-dependent loss of type H vessels and bone mass. According to our findings above, we reasoned that enhancing the expression of YBX1 may be a potential therapeutic strategy to alleviate ovariectomy-induced type H vessel richness reduction and bone mass loss. Our previous studies showed that some small molecular compounds that are naturally anti-inflammatory and antioxidative, including theaflavin 3-gallate (TF2A), Eriocitrin, Sciadopitysin, Bilobetin, and Isoginkgetin, target YBX1 by binding to its pocket-like structure (Appendix Fig S6 A-D and Fig 6 A)(Xiao *et al*, 2020). A cell proliferation assay was performed in endothelial cells to analyze the effect of 5 natural compounds on HUVECs cell viability. The results showed that TF2A and Sciadopitysin enhanced HUVECs cell viability, while Isoginkgetin significantly inhibited cell activity. And there is no detectable change in Eriocitrin and Bilobetin treatment (Appendix Fig S6 E). A tube formation assay with matrigel revealed that sciadopitysin treatment resulted in the substantial elevation of total branching points relative to vehicle control, while TF2A and Eriocitrin treatment was not altered (Appendix Fig S6 F and G). In addition, RT-qPCR analysis revealed that sciadopitysin treatment resulted in modestly higher transcript levels of BMP4, CD31, EMCN, and VEGFA, but displayed no effect on YBX1 transcript levels. Likewise, there was no detectable change in TF2A and Eriocitrin treatment (Appendix Fig S6 H). To further confirm the above findings, similar experiments were performed in HUVECs with different concentrations of sciadopitysin. We also discovered that sciadopitysin treatment of HUVECs increased endothelial cell migration (Appendix Fig S6 I and J), tube formation (Appendix Fig S6 K and L), and cell viability in a concentration-dependent manner (Fig 6 B). Moreover, western blotting results showed that treatment of endothelial cells isolated from primary bone with sciadopitysin led to upregulation of YBX1 and its target genes CD31, EMCN, and BMP4 expression (Fig 6 C). It is consistent with the result that sciadopitysin enhances Ybx1 stability by blocking the ubiquitination-mediated protein degradation pathway(Xiao *et al*., 2023). Next, we tested whether pharmacological administration of sciadopitysin ameliorated the reduction of CD31^hi^Emcn^hi^ endothelial cells and osteogenesis in ovariectomy mice. With a diameter of 20 to 80 nm and the ability to avoid immune recognition and clearance, biocompatible, biodegradable nanoparticles (NPs) exhibit excellent properties for carrying candidate drugs(Fish *et al*, 2021; Tunçay *et al*, 2000). So, we construct a nanocarrier by conjugating CD31 antibody (Alexa Fluor® 488 Conjugate) to NH2-polyethylene glycol (PEG) modified on PLGA poly (lactic-co-glycolic acid)-NH2-PEG-PLGA NPs to target endothelial cells (Appendix Fig S7 A). To directly assess the efficiency and specificity of CD31-NH2-PEG-PLGA NPs delivering sciadopitysin into endothelial cells in vitro, HUVECs were incubated with CD31-labeled NPs for 12 h and then stained with DAPI. We discovered a significant accumulation of green fluorescent NPs in HUVECs, indicating that CD31-labeled NPs could efficiently transport sciadopitysin to endothelial cells (Appendix Fig S7 B). We next modified CD31-labeled NPs with Cy7 dye and sought to analyze the biodistribution of the NPs. As shown, the Cy7 signal was readily detected in the heart, lung, spleen, liver, kidneys, and hind limbs, which validated the efficient distribution of CD31-labeled NPs to the skeletal and non-skeletal organs (537774v1).

**Fig 6.**
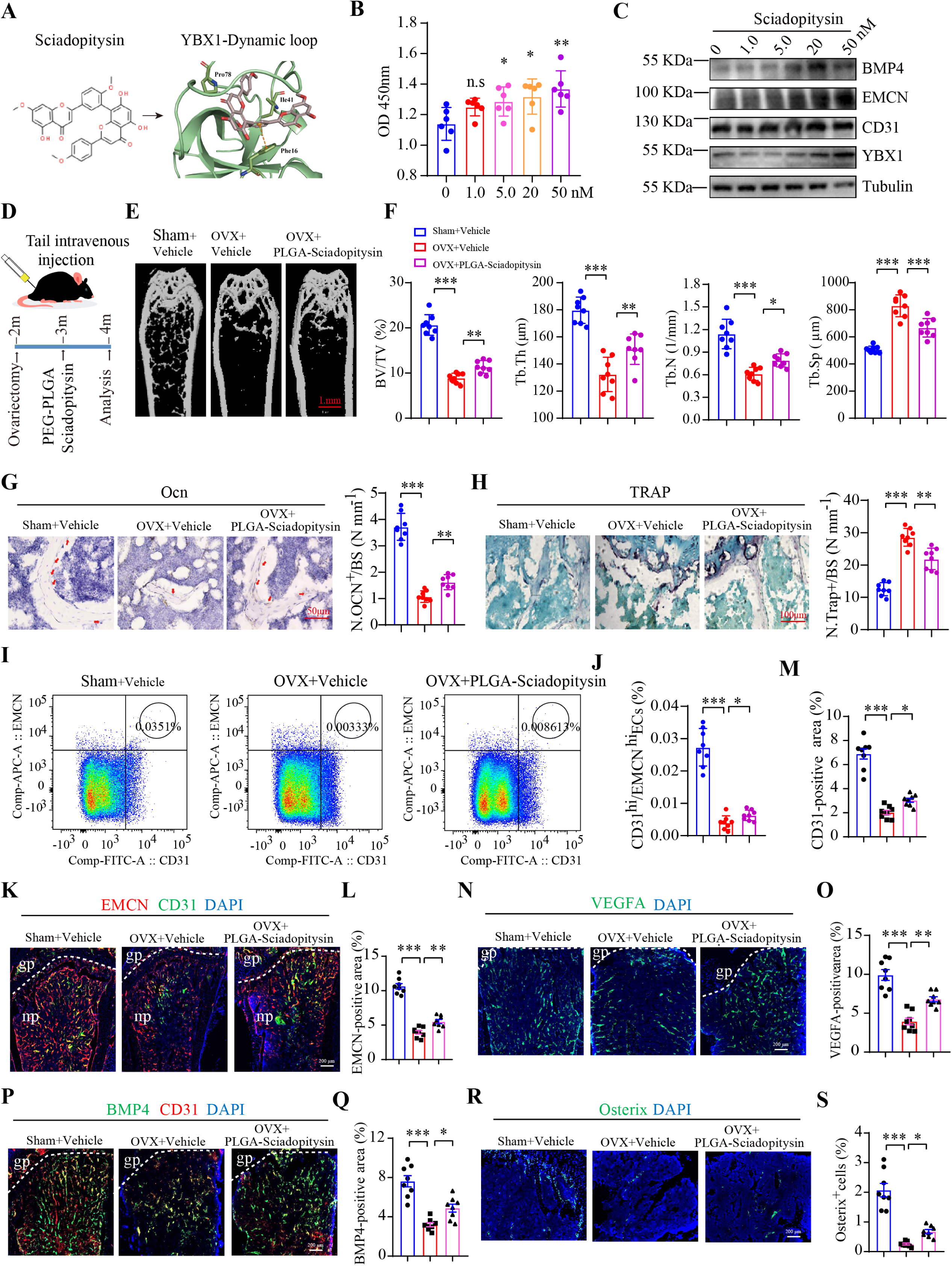
PEG-PLGA nanoparticles carrying sciadopitysin enhance angiogenesis-dependent bone formation in ovariectomy mice. A. The molecular structure of sciadopitysin (left) and the optimized binding modes with lowest binding energy generated by AutoDock Vina and key residues for interaction between sciadopitysin and mouse YBX1 (right). B. CCK8 assay analysis of the HUVECs viability after treating with different concentrations of sciadopitysin. C. Western blotting analysis of BMP4, EMCN, CD31 and YBX1 protein levels in HUVECs treated with different concentrations of sciadopitysin. D. Schematic diagram of treating OVX mice with sciadopitysin. E-F. Representative µCT images (E) and quantitative µCT analysis (F) of trabecular bone microarchitecture of sham, OVX, and OVX mice injected with PEG-PLGA nanoparticles carrying sciadopitysin (CD31 modified). Scale bar, 1mm. G. Immunohistochemical staining and quantification of Ocn^+^ cells (brown) in femora of sham, OVX, and OVX mice injected with PEG-PLGA nanoparticles carrying sciadopitysin (CD31 modified). Scale bar, 50 μm. H. Representative images of TRAP staining and quantification of TRAP^+^ cells in trabecular bone surfaces of sham, OVX, and OVX mice injected with PEG-PLGA nanoparticles carrying sciadopitysin (CD31 modified). Scale bar, 100 μm. I-J. FACS analysis dot plot (I) and quantification (J) of CD31^hi^EMCN^hi^ ECs from sham, OVX, and OVX mice injected with PEG-PLGA nanoparticles carrying sciadopitysin (CD31 modified). K-M. Representative images and quantitation of EMCN (red) and CD31 (green) immunostaining in tibia of sham, OVX, and OVX mice injected with PEG-PLGA nanoparticles carrying sciadopitysin (CD31 modified). Scale bar, 200 μm. N-O. Representative images and quantitation of VEGFA (green) immunostaining in tibia of sham, OVX, and OVX mice injected with PEG-PLGA nanoparticles carrying sciadopitysin (CD31 modified). Scale bar, 200 μm. P-Q. Representative images and quantitation of BMP4 (green) and CD31 (red) immunostaining in tibia of sham, OVX, and OVX mice injected with PEG-PLGA nanoparticles carrying sciadopitysin (CD31 modified). Scale bar, 200 μm. R-S. Representative images and quantitation of Osterix^+^ (green) immunostaining in tibia of sham, OVX, and OVX mice injected with PEG-PLGA nanoparticles carrying sciadopitysin (CD31 modified). Scale bar, 200 μm. n = 8 mice in each group. Data are shown as the mean ± SEM. *, P < 0.05; **, P < 0.01; ***, P < 0.001 by one-way ANOVA. Tb. BV/TV, trabecular bone volume per tissue volume; Tb. N, trabecular number; Tb. Sp, trabecular separation; Tb. Th, trabecular thickness.

Next, CD31-labeled NPs were injected in ovariectomy mice at day 30 post surgery (10 mg/kg) by tail vein injection three times a week for consecutive 30 days. As predicted, OVX mice treated with CD31-labeled NPs had significantly increased bone mass, osteoblast number, and osteoprogenitor cells (Fig 6 E-G and R-S). Meanwhile, the osteoclast number was significantly reduced compared with OVX control mice (Fig 6 H). Moreover, flow cytometric quantification at 30 days post-irradiation showed the amounts of type H endothelial cells sub-population were significantly increased in comparison with OVX control mice (Fig 6 I and J). Furthermore, 30 days of CD31-labeled NPs treatment led to substantial expansion of CD31^hi^EMCN^hi^ endothelial cells in OVX mice (Fig 6 K-M). Likewise, VEGFA and BMP4 fluorescent intensities were significantly increased (Fig 6 N-Q). Taken together, these results give proof of the principle that CD31-labeled sciadopitysin NPs might have clinical utility to ameliorate disorders of low bone mass like postmenopausal osteoporosis. Next, to determine the therapeutic effects of sciadopitysin on endothelium formation and bone formation in aging mice, CD31-labeled NPs were injected into 20-month-old mice (10mg/kg) by tail vein injection three times a week for consecutive 30 days (Fig 7 A). Flow cytometry analysis showed that the administration of CD31-labeled NPs substantially countered age-induced CD31^hi^EMCN^hi^ endothelium loss (Fig 7 B and C). Likewise, CD31^hi^Emcn^hi^ endothelial cells density (Fig 7 D-F), VEGFA (Fig 7 G and H) and BMP4 fluorescent intensities (Fig 7 I and J), and osterix^+^ osteoprogenitors cells were similarly improved in comparison with control mice (Fig 7 K and L).

**Fig 7.**
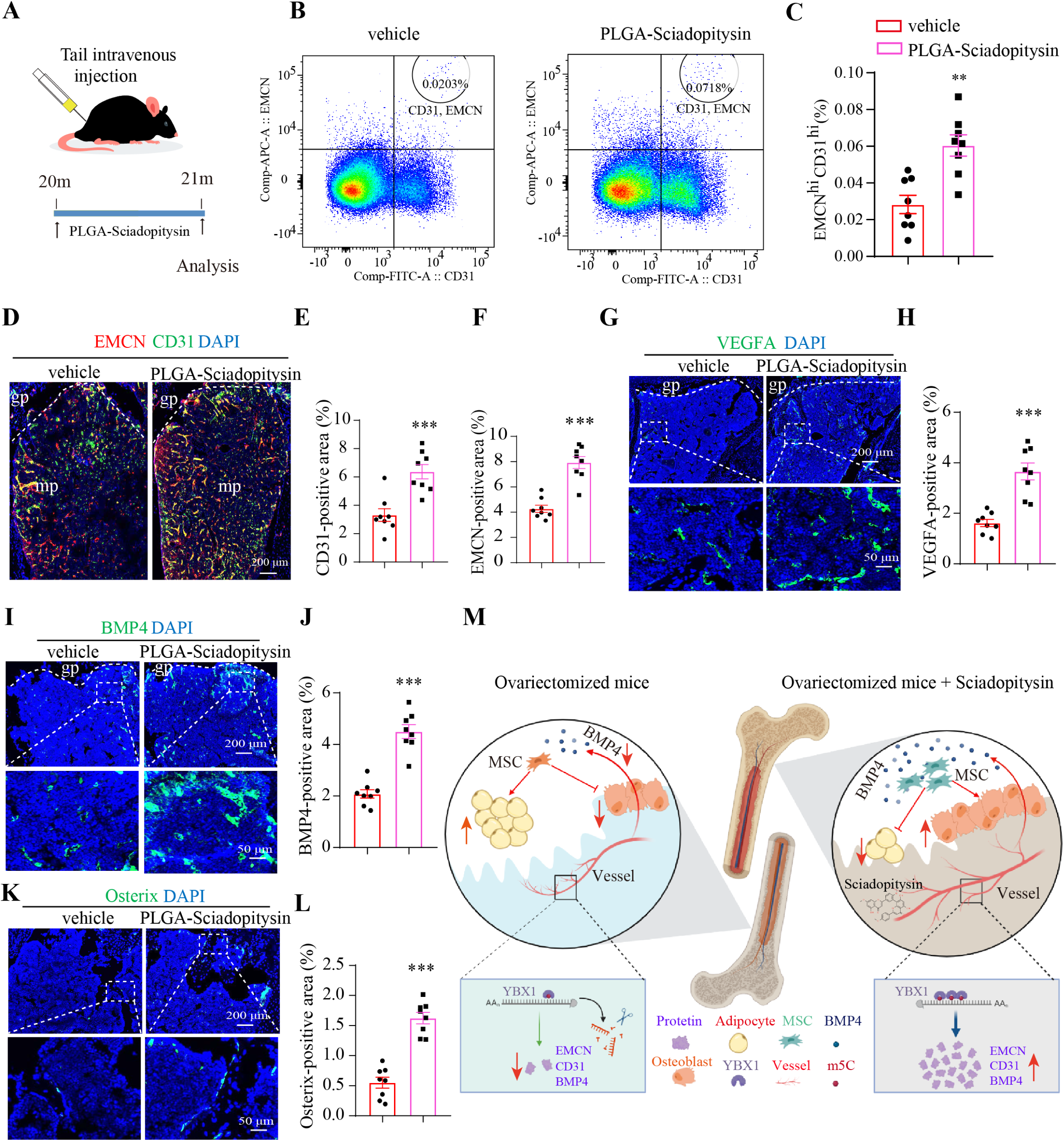
PEG-PLGA nanoparticles carrying sciadopitysin enhance angiogenesis-dependent bone formation in aged mice. A. Schematic diagram of treating aged mice with sciadopitysin. B-C. FACS analysis dot plot (B) and quantification (C) of CD31^hi^EMCN^hi^ ECs from aged mice injected with vehicle (PEG-PLGA nanoparticles) and PEG-PLGA nanoparticles carrying sciadopitysin (CD31 modified). D-F. Representative images and quantitation of EMCN (red) and CD31 (green) immunostaining in tibia of aged mice injected with vehicle and PEG-PLGA nanoparticles carrying sciadopitysin (CD31 modified). Scale bar, 200 μm. G-H. Representative images and quantitation of VEGFA (green) immunostaining in tibia of aged mice injected with vehicle and PEG-PLGA nanoparticles carrying sciadopitysin (CD31 modified). Scale bar, 200 μm. I-J. Representative images and quantitation of BMP4 (green) immunostaining in tibia of aged mice injected with vehicle and PEG-PLGA nanoparticles carrying sciadopitysin (CD31 modified). Scale bar, 200 μm. K-L. Representative images and quantitation of Osterix^+^ (green) immunostaining in tibia of aged mice injected with vehicle and PEG-PLGA nanoparticles carrying sciadopitysin (CD31 modified). Scale bar, 200 μm. M. Schematic of the reduced YBX1 that impaired type H vessel formation by influencing BMP4, CD31 and EMCN stability in an m5C-dependent manner and then inhibited osteogenic differentiation and promoted adipogenetic differentiation of BMSCs in OVX mice, which were restored by treating with sciadopitysin coated by PEG-PLGA nanoparticles. n = 8 mice in each group. Data are shown as the mean ± SEM. **, P < 0.01; ***, P < 0.001 by student’s t test.

Taken together, these results suggested that elevating the expression of Ybx1 by intravenous administration of sciadopitysin promoted CD31^hi^EMCN^hi^ vessel formation and stimulated new bone formation in both aged and OVX osteoporosis mouse models (Fig 7 M).

## Discussion

RNA binding proteins (RBPs) are a huge and complicated group that is essential to post-transcriptional gene regulation. Lesser known, but not less important, the RBPs can also regulate the formation of vessel in tissues(Smith & Costa, 2022). In the present study, we are the first to elucidate that RBP-Ybx1 promotes Type H vessels - dependent bone formation by regulating CD31 and BMP4 stability in an m5C-dependent manner. As we know, Ybx1 is a multi-function factor that involves the processes of mRNA transcription, stability, splicing, protein packaging, and translation(Lyabin *et al*, 2014; Wu *et al*, 2023). Our a previous study revealed that Ybx1, as an alternative splicing of pre-mRNA, regulates BMSCs senescence and differentiation during aging(Xiao *et al*., 2023). In fact, alternative splicing of pre-mRNA is an important post-transcriptional way that governs the process of angiogenesis by generating alternative spliced isoforms of key pro-angiogenesis genes including VEGF-A, VEGFR1, VEGFR2, NRP-1, FGFRs, Vasohibin-1, Vasohibin-2, HIF-1α, Angiopoietin-1 and Angiopoietin-2(Bowler & Oltean, 2019). However, in our study, we didn’t find any significant alterations of alternative spliced isoforms of these candidate pro-angiogenesis genes. Nevertheless, we cannot exclude the possibility that Ybx1 regulates Type H vessels formation by affecting pre-mRNA alternative splicing of other angiogenesis-related genes.

Another important finding in the present study was the recognition of the communication between Type H vessels and BMSCs differentiation. Previous studies were focused on the coupling of Type H vessels and osteoprogenitors but ignored the coupling of Type H vessels and BMSCs. Like the osteoprogenitors, BMSCs are also surrounded with vessels in bone tissues (figure 5 A)(Tuckermann & Adams, 2021). In the case of bone formation, BMSCs are multilineage potential stem cells and are the source of new osteoblasts and their progenitors, which are essential for bone homeostasis and fracture healing(Pittenger *et al*, 1999). Studies supported that the age-related switch between osteoblast and adipocyte differentiation of BMSCs is a major factor for age-related bone loss(Guilak *et al*, 2009; Li *et al*, 2015). Here we demonstrated that deletion of Ybx1 in endothelial cells decreased the expression of endothelial cell-derived Bmp4 and then disrupted the switch between adipogenic differentiation and osteogenic differentiation of BMSCs. Our results further demonstrate that endothelial cells are often a source of paracrine molecular signals and control the behaviour of other cell types in the surrounding tissue by secreting endothelial cell-derived instructive factors including Bmp4, Tgfb2, Nog, Fgf1, Fgf8, Wnt1, Wnt3a, Wnt10b, Dkk1 and Pgf(Langen *et al*., 2017; Ramasamy *et al*, 2014). We also propose that type H endothelial cells mediate local proliferation of BMSCs and provide niche signals for BMSCs.

There is no doubt that BMP4 has been implicated in the induction of osteoblast differentiation during embryonic skeletogenesis and fracture healing(Abe *et al*, 2000; Langen *et al*., 2017; Pan *et al*, 2023). Meanwhile, BMP4-induced commitment to the osteoblastic lineage is a prerequisite for osteoclast development and BMP4 depletion or treatment with a BMP4 signaling inhibitor diminished osteoclast differentiation(Abe *et al*., 2000; Zuo *et al*, 2021). As such, it is reasonable to assume that osteoclastogenesis is impaired in in the Ybx1^iΔEC^ mice. However, as we show in the following manuscript, deletion of Ybx1 in endothelial cells instead promoted osteoclastogenesis. TRAP stain analysis in Ybx1^iΔEC^ mice femur showed that a mass of osteoclasts was enriched on the bone surface (Fig 2 P and Q). It suggested that abnormally activated osteoclastic differentiation may be considered as another cause for inducing of Ybx1 deletion-induced bone loss. This result also strongly suggests the existence of other factors that promote osteoclast differentiation. Excitingly, we found the TNFα mRNA levels, a crucial differentiation factor for osteoclasts activity(Azuma *et al*, 2000), was significantly increased in shYbx1 samples by analyzing the RNA-seq data. The result was further confirmed by measuring the TNFα levels in the medium from Ybx1^iΔEC^ mice primary bone morrow (BM)-derived endothelial cells (Appendix Fig S5 A-C). In addition, the promotional effect of Ybx1^iΔEC^ conditioned medium on osteoclastic differentiation was counteracted by TNFα neutralizing antibody (Appendix Fig S5 F and G). The transcription factor analysis experiments indicated that Ybx1 may regulate the expression of IRF1 and then indirectly regulate TNFα levels (Appendix Fig S5 D and E). However, the exact molecular mechanism underlying the different role of Ybx1 in osteoclastic differentiation remain to be elucidated.

Poly (lactic-co-glycolic acid) (PLGA) is one of biodegradable polymeric nanoparticles (NPs) with well ustained-release properties, low toxicity, and biocompatibility with tissue and cells and had been widely applied in drug delivery systems for the treatment of many diseases(Sadat Tabatabaei Mirakabad *et al*, 2014). Sciadopitysin is a natural compound isolated from the traditional Chinese herbal agent with well anti-oxidant, anti-osteoclastogenesis and anti-Alzheimer’s disease(Cao *et al*, 2017; Gu *et al*, 2013; Suh *et al*, 2013). Our previous study supported that sciadopitysin enhanced the protein expression of Ybx1 by blocking the ubiquitinated degradation pathway. However, its poor water solubility remains clinical challenges. In this study, we examined the possibility of loading sciadopitysin into CD31-modified NPs and tested the effectiveness of treatment for the decline of type H vessels and bone mass in OVX and aging mice. In our study, we found that sciadopitysin is selectively delivered to the vasculature by CD31-modified NPs, and that the defective bone angiogenesis and osteogenesis in OVX and elderly mice were successfully reversed after targeted administration of CD31-modified sciadopitysin NPs. We believe this work offers an avenue to increase the clinical utility of sciadopitysin in the treatment of osteoporosis diseases.

In conclusion, we revealed that YBX1 acts as an RNA-binding protein that promotes Type H vessels dependent bone formation by coupling with BMSCs differentiation. The CD31-modified sciadopitysin NPs can be used to boost type H vessel formation and osteogenesis in aged and OVX mice. Our findings presented here set the groundwork for novel therapeutic approaches in the management of osteoporotic disease states by boosting angiogenesis dependent osteogenesis.

## Data availability

All the data that support the findings of this study are available from the corresponding author upon reasonable request.

## Acknowledgments

This work was supported by grants from National Natural Science Foundation of China (Grant No.81900810)

## Author contributions

Yu-Jue Li generated data and drafted the manuscript; Qi Guo analyzed results and revised the manuscript; Wen-Feng Xiao supervised the experiments and co-wrote the manuscript; Ye Xiao designed the experiments, curated data, acquisited funding and revised the manuscript.

## Figure Legend

**Appendix Figure 1.**
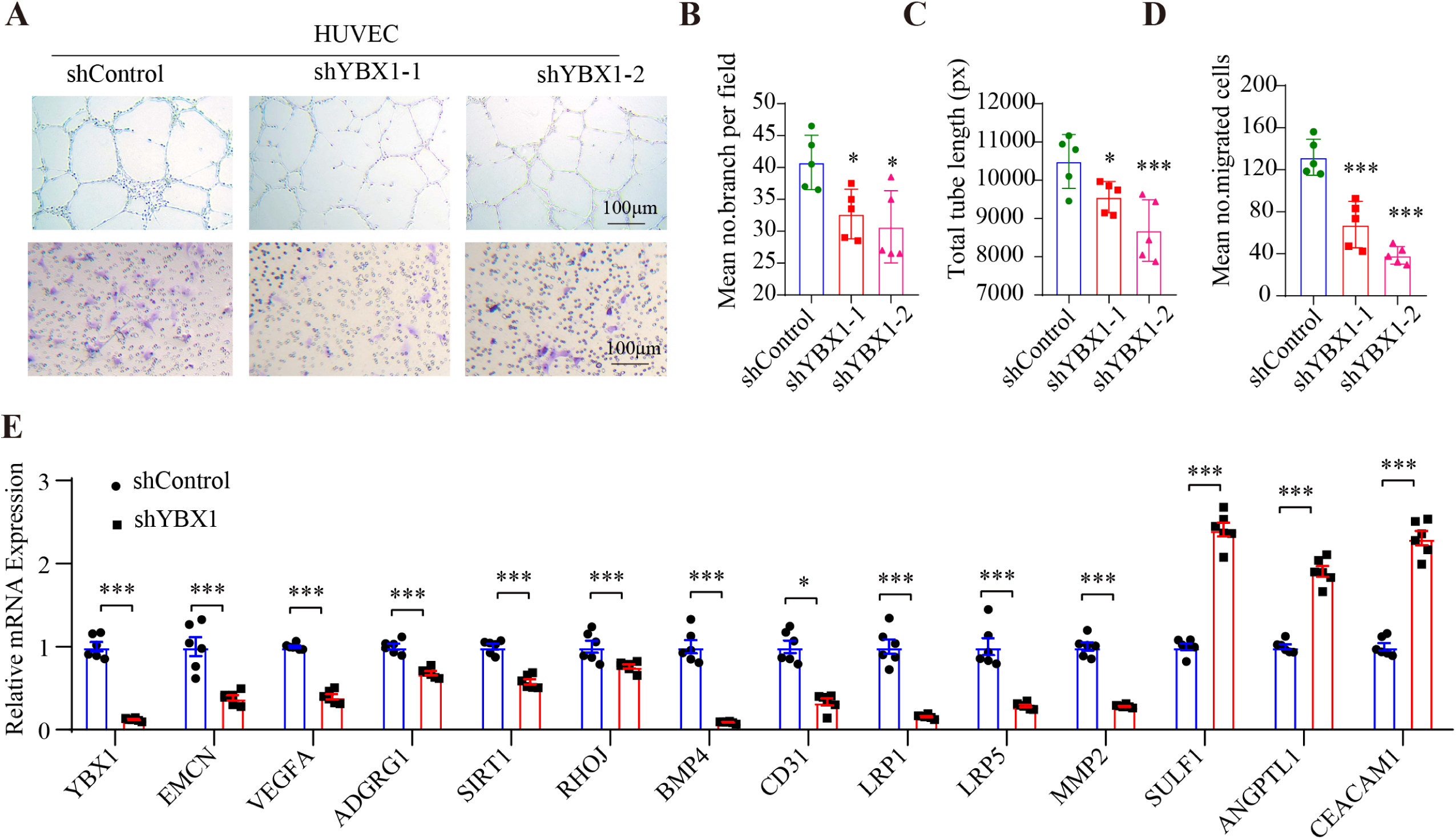
Slicing of YBX1 damaged angiogenesis of HUVECs. A-D. Representative images (A) and relative quantification (B-D) of tube branch numbers of a Matrigel tube formation assay (up panel) and a transwell migration assay (bottom panel). E. RT-qPCR analysis of the expression of angiogenesis-associated genes in HUVECs infected with shYBX1 adenovirus or control adenovirus. n = 3 independent experiments. Data are shown as the mean ± SEM. *, P < 0.05; **, P < 0.01; ***, P < 0.001 by one-way ANOVA (B-D) or Student’s t test (E).

**Appendix Figure 2.**
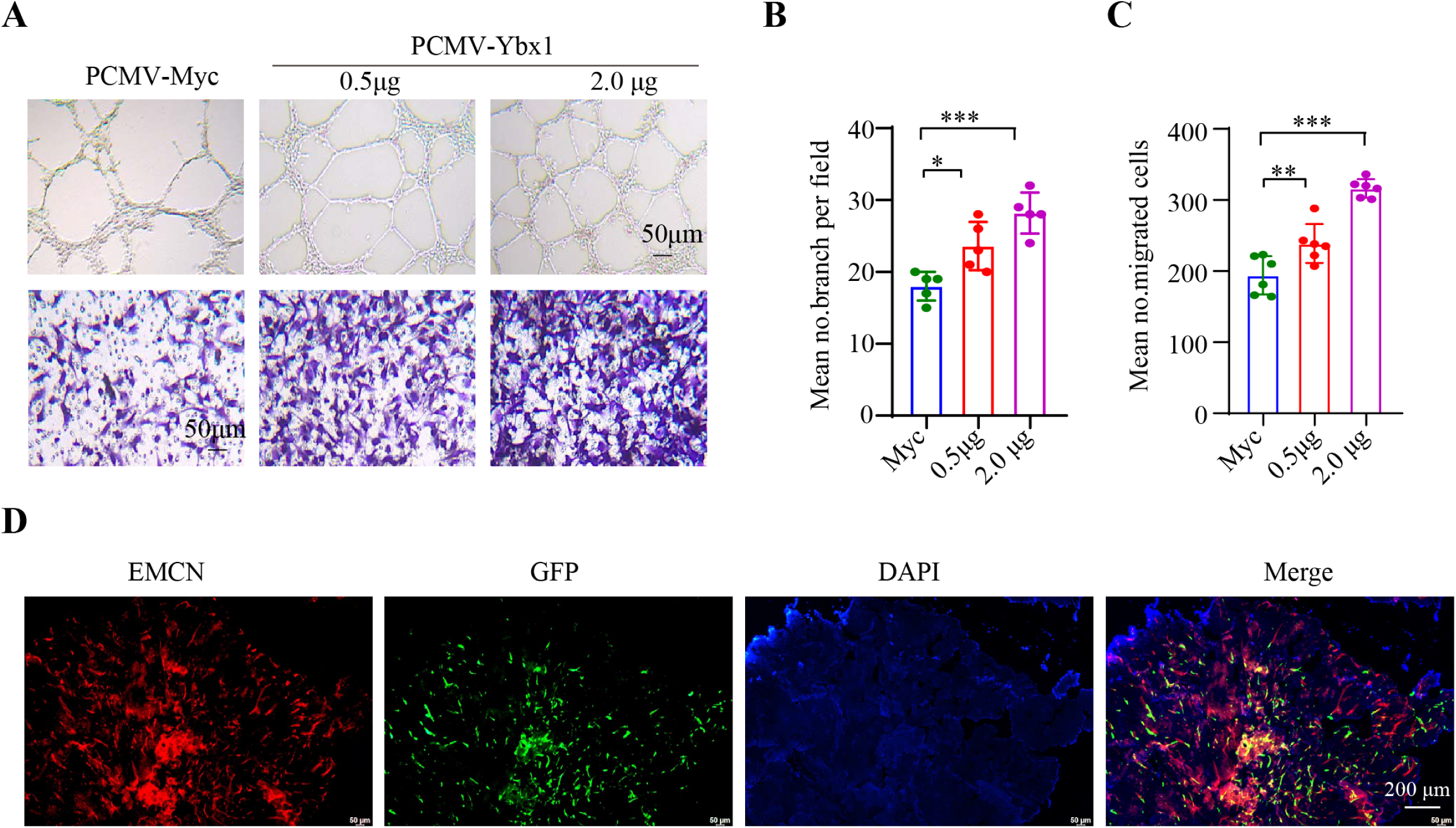
Over-expression of YBX1 enhanced angiogenesis of HUVECs. A-C. Representative images (A) and relative quantification (B-C) of tube branch numbers of a Matrigel tube formation assay (up panel) and a transwell migration assay (bottom panel). n = 3 independent experiments. D. Representative images of EMCN (red) and GFP (green) immunostaining in femora infected with rAAV2/9-Tie-Ybx1 or rAAV 2/9. Data are shown as the mean ± SEM. *, P < 0.05; **, P < 0.01; ***, P < 0.001 by one-way ANOVA.

**Appendix Figure 3.**
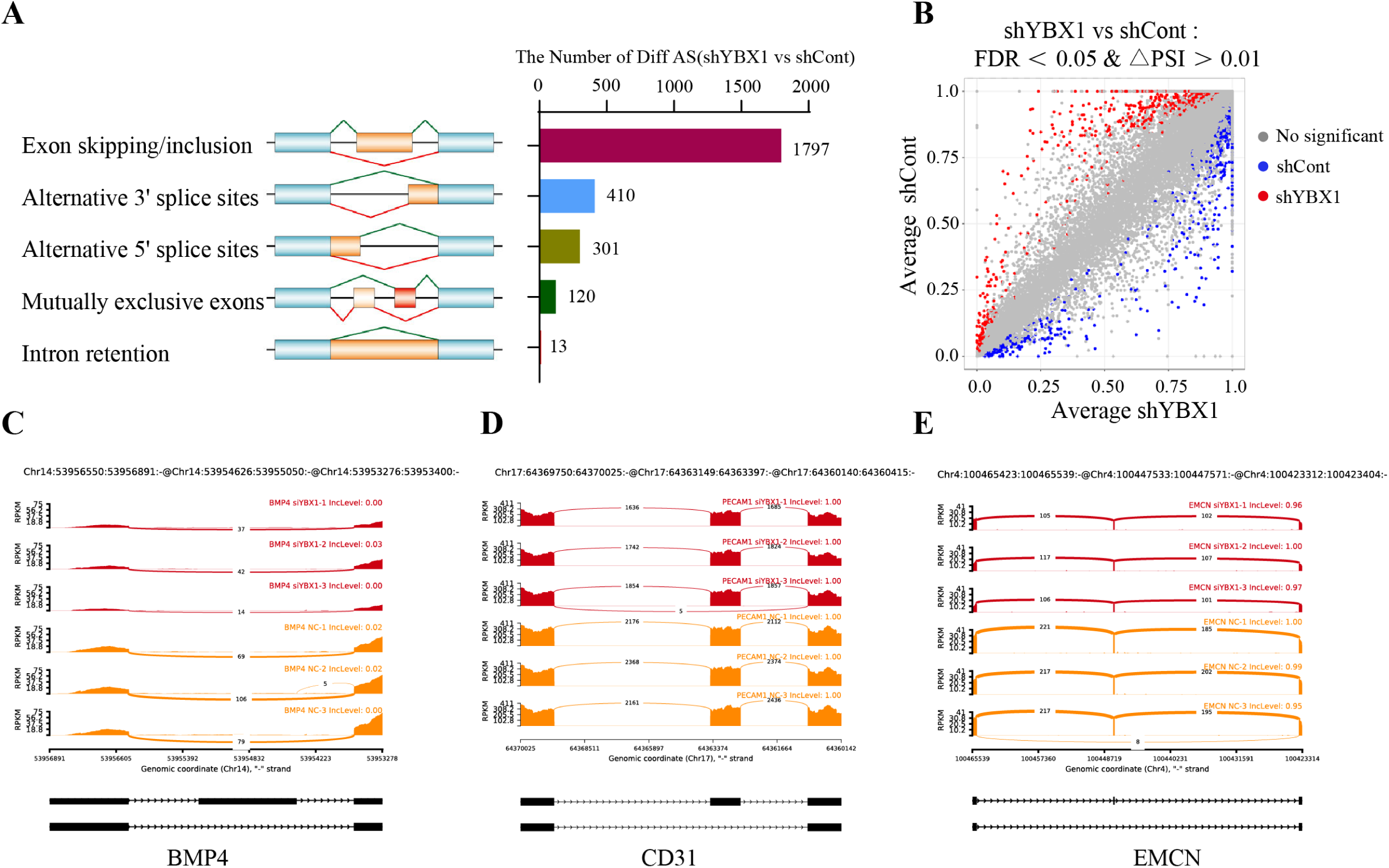
Deletion of YBX1 in endothelial cells had no effect on BMP4, CD31, and EMCN pre-mRNA alternative splicing. A. Schematic of pre-mRNA alternative splicing and the number of differentially spliced events in HUVECs infected with shYBX1 adenovirus or control adenovirus. B. Scatterplot of exons promoted (red circles) and repressed (blue circles) in HUVECs infected with shYBX1 adenovirus or control adenovirus. C-E. Sashimi plots of BMP4, CD31 and EMCN in HUVECs infected with shYBX1 adenovirus or control adenovirus. n = 3 independent experiments.

**Appendix Figure 4.**
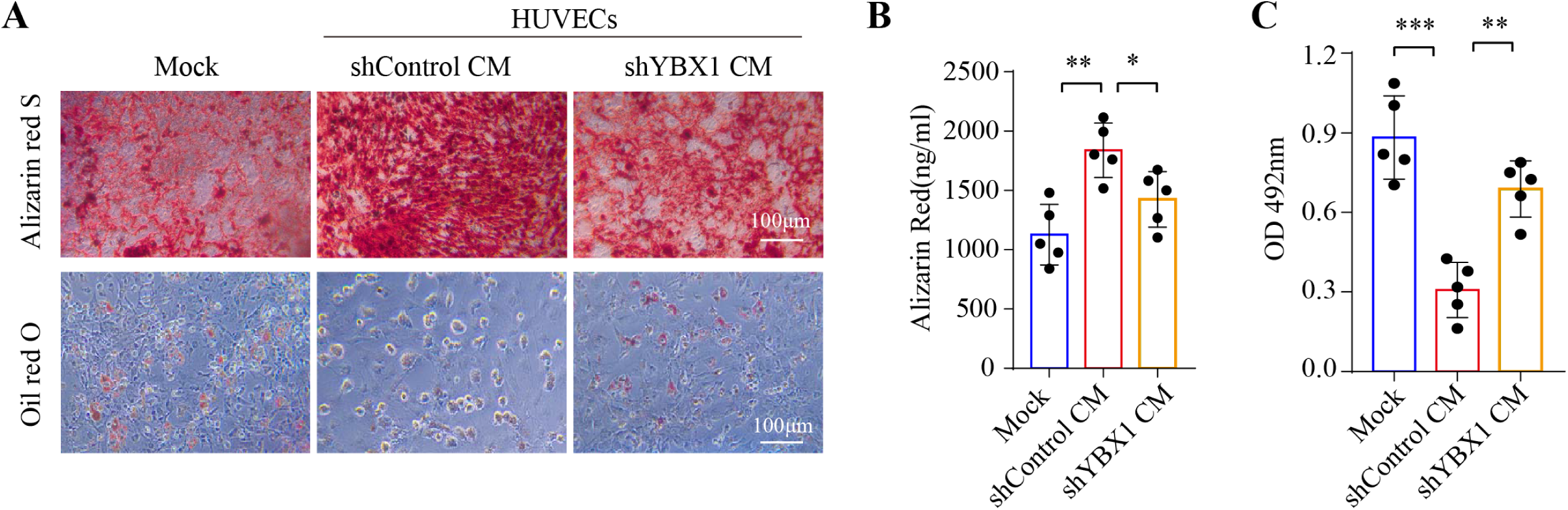
HUVECs medium have an effect on osteogenic differentiation and adipogenic differentiation of BMSCs. A. Representative images of Alizarin Red staining (up panel) and Oil Red O staining (bottom panel) in BMSCs treated with conditioned medium from HUVECs infected with shYBX1 adenovirus or control adenovirus. B-C. Quantification of calcium mineralization based on Alizarin Red staining (B) and quantification of Oil Red O based on Oil Red O staining (C) in BMSCs treated with conditioned medium. n = 3 independent experiments. Data are shown as the mean ± SEM. *, P < 0.05; **, P < 0.01; ***, P < 0.001 by one-way ANOVA.

**Appendix Figure 5.**
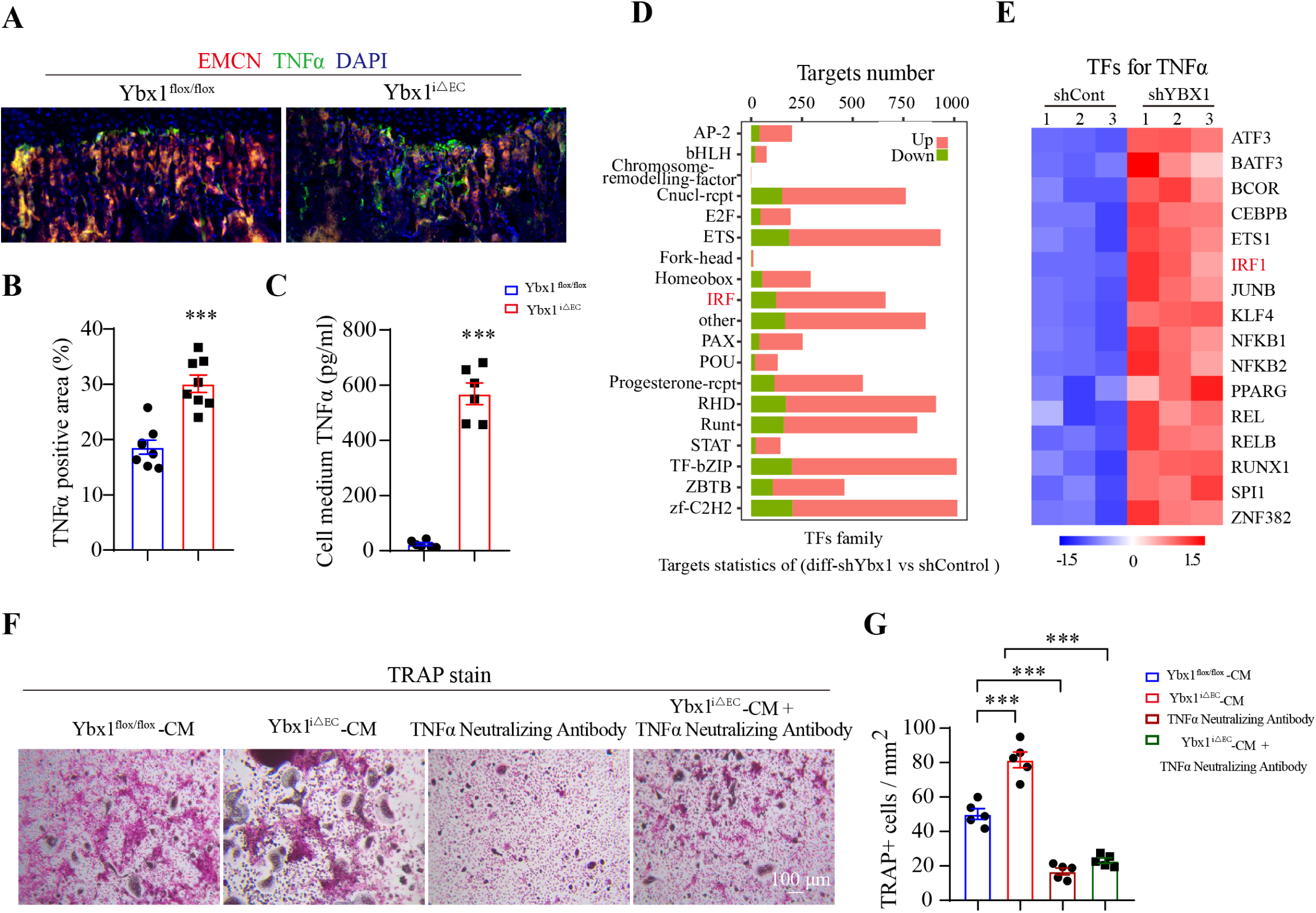
Deletion of YBX1 in endothelial cells activated TNFα expression. A-B. Representative images and quantitation of EMCN (red) and TNFα (green) immunostaining in femora from endothelial-specific YBX1 knockout mice (Ybx1^iΔEC^) and their littermate controls (Ybx1^flox/flox^). C. ELISA analysis of TNFα levels in cell medium from isolated Ybx1^iΔEC^ and Ybx1^flox/flox^ endothelial cells. D. The altered transcription factors family and the number of target genes of transcription factors in endothelial cells infected with shYBX1 adenovirus or control adenovirus. E. A heat-map of altered transcription factors involves regulating TNFα between endothelial cells infected with shYBX1 adenovirus or control adenovirus. F-G. Representative images of TRAP staining (F) and quantification of staining of osteoclasts treating with conditioned medium from HUVECs infected with shYBX1 adenovirus or control adenovirus or TNFα neutralizing antibody (G). n = 3 independent experiments. Data are shown as the mean ± SEM. ***, P < 0.001 by Student’s t test (B and C) and two-way ANOVA (G).

**Appendix Figure 6.**
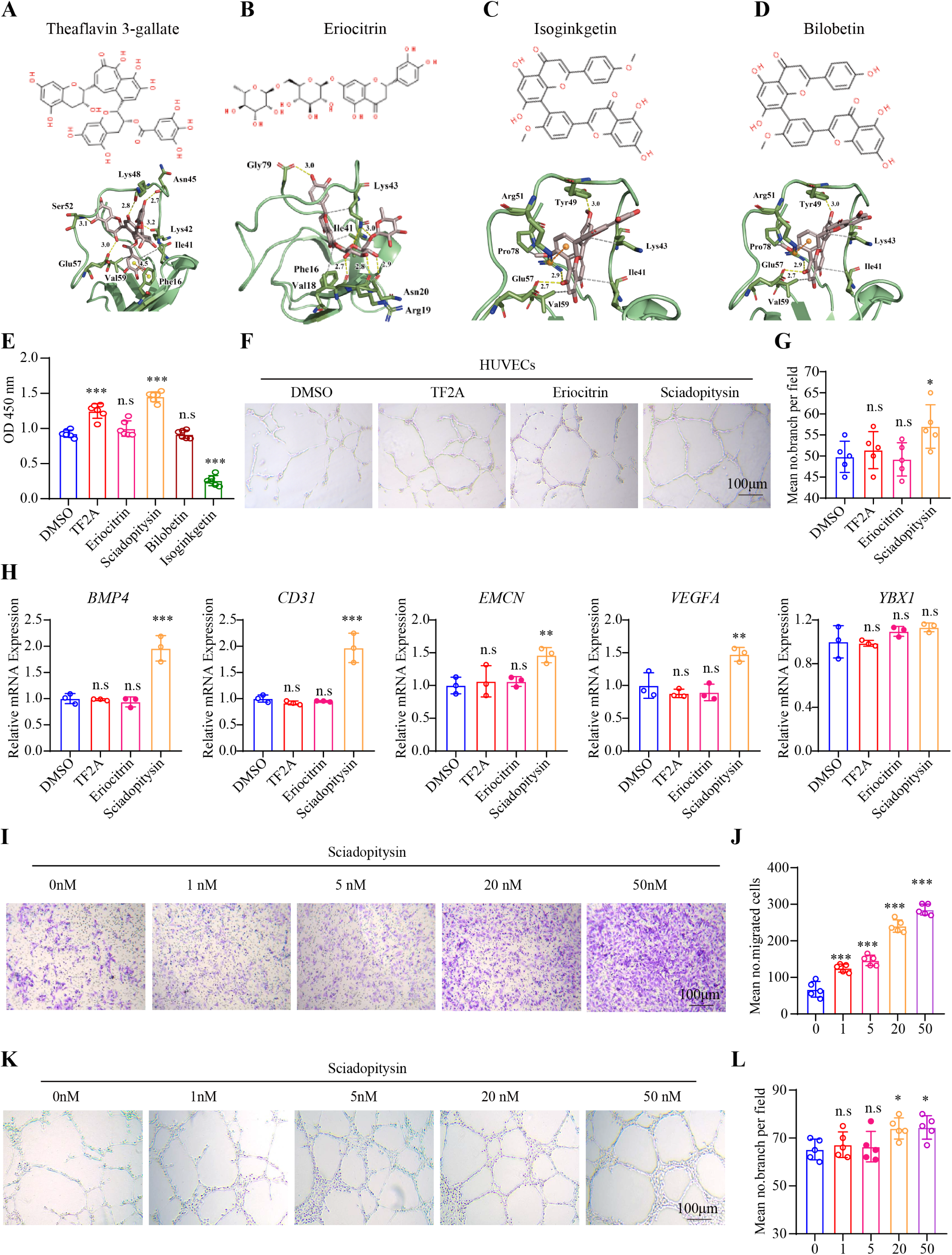
Screening for natural small molecular compounds that activate YBX1 expression. A-D. The Theaflavin-3-gallate (TF2A), Eriocitrin, Isoginkgetin, and Bilobetin structure (upper panel) and the key residues for interaction between them and YBX1 (bottom panel). E. CCK8 assay analysis of the HUVECs viability after treating with (TF2A), Eriocitrin, Sciadopitysin, Bilobetin, and Isoginkgetin. F-G. Representative images (F) and relative quantification (G) of tube branch numbers of HUVECs treated with TF2A, Eriocitrin and Sciadopitysin. H. BMP4, CD31, EMCN, VEGFA and YBX1 mRNA expression in HUVECs treated with TF2A, Eriocitrin and Sciadopitysin. I-J. Representative images (I) and relative quantification (J) of migrating cells treated with different concentrations of sciadopitysin. K-L. Representative images (K) and relative quantification (L) of tube branch numbers of HUVECs treated with different concentrations of sciadopitysin. n = 3 independent experiments. Data are shown as the mean ± SEM. *, P < 0.05; **, P < 0.01; ***, P < 0.001 by one-way ANOVA.

**Appendix Figure 7.**
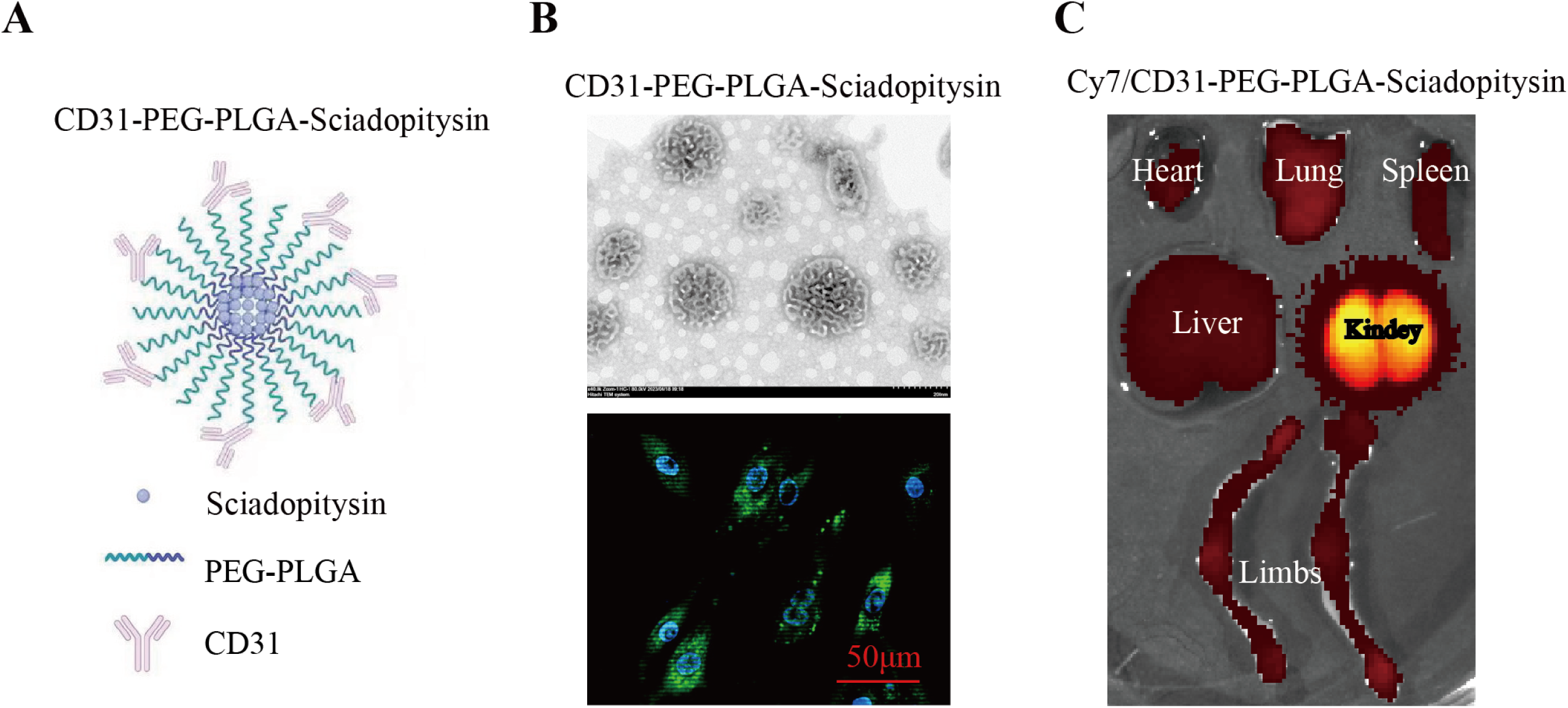
PEG-PLGA-sciadopitysin nanoparticles effectively target endothelial cells. A. Schematic of CD31-modified PEG-PLGA-sciadopitysin nanoparticles. B. Representative image of scanning electron microscopy of CD31-modified PEG-PLGA-sciadopitysin nanoparticles (up panel) and the nanoparticles were transferred into HUVECs (bottom panel). C. Representative ex vivo fluorescence images of heart, lung, spleen, liver, kidneys, and hind limbs of 8-week-old mice, which were injected with Cy7/CD31-modified PEG-PLGA-sciadopitysin nanoparticles by tail vein and sacrificed 24 h post injection.

---

**Table 1.**
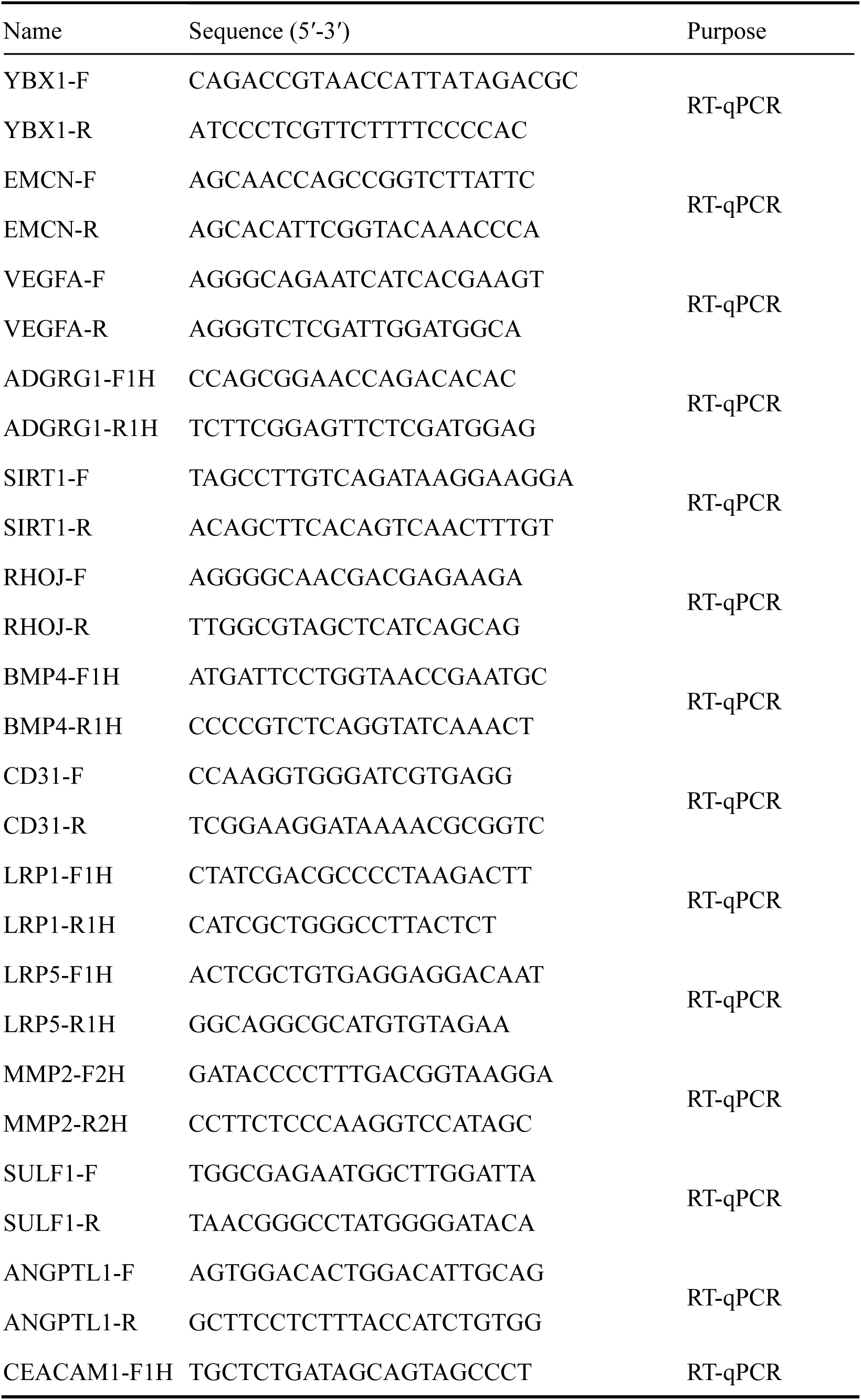
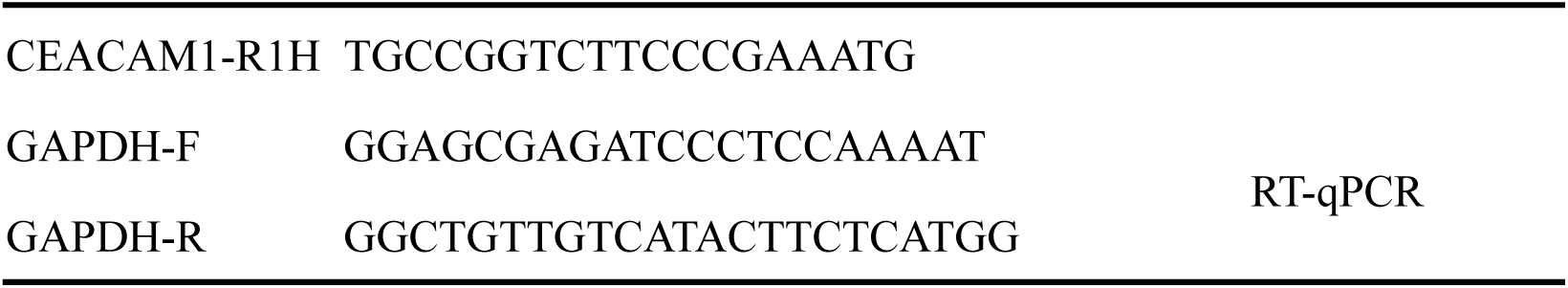
The primer sequences

## References

Abe E, Yamamoto M, Taguchi Y, Lecka-Czernik B, O’Brien CA, Economides AN, Stahl N, Jilka RL, Manolagas SC (2000) Essential requirement of BMPs-2/4 for both osteoblast and osteoclast formation in murine bone marrow cultures from adult mice: antagonism by noggin. J Bone Miner Res 15: 663–673

Azuma Y, Kaji K, Katogi R, Takeshita S, Kudo A (2000) Tumor necrosis factor-alpha induces differentiation of and bone resorption by osteoclasts. J Biol Chem 275: 4858–4864

Baccin C, Al-Sabah J, Velten L, Helbling PM, Grünschläger F, Hernández-Malmierca P, Nombela-Arrieta C, Steinmetz LM, Trumpp A, Haas S (2020) Combined single-cell and spatial transcriptomics reveal the molecular, cellular and spatial bone marrow niche organization. Nat Cell Biol 22: 38–48

Bowler E, Oltean S (2019) Alternative Splicing in Angiogenesis. Int J Mol Sci 20

Cao J, Lu Q, Liu N, Zhang YX, Wang J, Zhang M, Wang HB, Sun WC (2017) Sciadopitysin suppresses RANKL-mediated osteoclastogenesis and prevents bone loss in LPS-treated mice. Int Immunopharmacol 49: 109–117

Chang SH, Hla T (2011) Gene regulation by RNA binding proteins and microRNAs in angiogenesis. Trends Mol Med 17: 650–658

Chen J, Li M, Liu AQ, Zheng CX, Bao LH, Chen K, Xu XL, Guan JT, Bai M, Zhou T et al (2020) Gli1(+) Cells Couple with Type H Vessels and Are Required for Type H Vessel Formation. Stem Cell Reports 15: 110–124

Chen X, Li A, Sun BF, Yang Y, Han YN, Yuan X, Chen RX, Wei WS, Liu Y, Gao CC et al (2019) 5-methylcytosine promotes pathogenesis of bladder cancer through stabilizing mRNAs. Nat Cell Biol 21: 978–990

Coles LS, Bartley MA, Bert A, Hunter J, Polyak S, Diamond P, Vadas MA, Goodall GJ (2004) A multi-protein complex containing cold shock domain (Y-box) and polypyrimidine tract binding proteins forms on the vascular endothelial growth factor mRNA. Potential role in mRNA stabilization. Eur J Biochem 271: 648–660

Coles LS, Diamond P, Lambrusco L, Hunter J, Burrows J, Vadas MA, Goodall GJ (2002) A novel mechanism of repression of the vascular endothelial growth factor promoter, by single strand DNA binding cold shock domain (Y-box) proteins in normoxic fibroblasts. Nucleic Acids Res 30: 4845–4854

Coles LS, Lambrusco L, Burrows J, Hunter J, Diamond P, Bert AG, Vadas MA, Goodall GJ (2005) Phosphorylation of cold shock domain/Y-box proteins by ERK2 and GSK3beta and repression of the human VEGF promoter. FEBS Lett 579: 5372–5378

Cook KB, Kazan H, Zuberi K, Morris Q, Hughes TR (2011) RBPDB: a database of RNA-binding specificities. Nucleic Acids Res 39: D301–308

Cui Z, Wu H, Xiao Y, Xu T, Jia J, Lin H, Lin R, Chen K, Lin Y, Li K et al (2022) Endothelial PDGF-BB/PDGFR-β signaling promotes osteoarthritis by enhancing angiogenesis-dependent abnormal subchondral bone formation. Bone Res 10: 58

El-Naggar AM, Veinotte CJ, Cheng H, Grunewald TG, Negri GL, Somasekharan SP, Corkery DP, Tirode F, Mathers J, Khan D et al (2015) Translational Activation of HIF1α by YB-1 Promotes Sarcoma Metastasis. Cancer Cell 27: 682–697

Ferrara N, Adamis AP (2016) Ten years of anti-vascular endothelial growth factor therapy. Nat Rev Drug Discov 15: 385–403

Fish MB, Banka AL, Braunreuther M, Fromen CA, Kelley WJ, Lee J, Adili R, Holinstat M, Eniola-Adefeso O (2021) Deformable microparticles for shuttling nanoparticles to the vascular wall. Sci Adv 7

Fu R, Lv WC, Xu Y, Gong MY, Chen XJ, Jiang N, Xu Y, Yao QQ, Di L, Lu T et al (2020) Endothelial ZEB1 promotes angiogenesis-dependent bone formation and reverses osteoporosis. Nat Commun 11: 460

Gao D, Niu Q, Gong Y, Guo Q, Zhang S, Wang Y, Liu S, Wang H, Svatek R, Rodriguez R et al (2021) Y-Box Binding Protein 1 Regulates Angiogenesis in Bladder Cancer via miR-29b-3p-VEGFA Pathway. J Oncol 2021: 9913015

Georgilis A, Klotz S, Hanley CJ, Herranz N, Weirich B, Morancho B, Leote AC, D’Artista L, Gallage S, Seehawer M et al (2018) PTBP1-Mediated Alternative Splicing Regulates the Inflammatory Secretome and the Pro-tumorigenic Effects of Senescent Cells. Cancer Cell 34: 85–102.e109

Gerstberger S, Hafner M, Tuschl T (2014) A census of human RNA-binding proteins. Nat Rev Genet 15: 829–845

Gu Q, Li Y, Chen Y, Yao P, Ou T (2013) Sciadopitysin: active component from Taxus chinensis for anti-Alzheimer’s disease. Nat Prod Res 27: 2157–2160

Guilak F, Cohen DM, Estes BT, Gimble JM, Liedtke W, Chen CS (2009) Control of stem cell fate by physical interactions with the extracellular matrix. Cell Stem Cell 5: 17–26

Huang J, Yin H, Rao SS, Xie PL, Cao X, Rao T, Liu SY, Wang ZX, Cao J, Hu Y et al (2018) Harmine enhances type H vessel formation and prevents bone loss in ovariectomized mice. Theranostics 8: 2435–2446

Jayavelu AK, Schnöder TM, Perner F, Herzog C, Meiler A, Krishnamoorthy G, Huber N, Mohr J, Edelmann-Stephan B, Austin R et al (2020) Splicing factor YBX1 mediates persistence of JAK2-mutated neoplasms. Nature 588: 157–163

Ji X, Kong J, Liebhaber SA (2011) An RNA-protein complex links enhanced nuclear 3’ processing with cytoplasmic mRNA stabilization. Embo j 30: 2622–2633

Kusumbe AP, Ramasamy SK, Adams RH (2014) Coupling of angiogenesis and osteogenesis by a specific vessel subtype in bone. Nature 507: 323–328

Langen UH, Pitulescu ME, Kim JM, Enriquez-Gasca R, Sivaraj KK, Kusumbe AP, Singh A, Di Russo J, Bixel MG, Zhou B et al (2017) Cell-matrix signals specify bone endothelial cells during developmental osteogenesis. Nat Cell Biol 19: 189–201

Lee YC, Cheng CJ, Bilen MA, Lu JF, Satcher RL, Yu-Lee LY, Gallick GE, Maity SN, Lin SH (2011) BMP4 promotes prostate tumor growth in bone through osteogenesis. Cancer Res 71: 5194–5203

Li CJ, Cheng P, Liang MK, Chen YS, Lu Q, Wang JY, Xia ZY, Zhou HD, Cao X, Xie H et al (2015) MicroRNA-188 regulates age-related switch between osteoblast and adipocyte differentiation. J Clin Invest 125: 1509–1522

Li CJ, Xiao Y, Sun YC, He WZ, Liu L, Huang M, He C, Huang M, Chen KX, Hou J et al (2021) Senescent immune cells release grancalcin to promote skeletal aging. Cell Metab 33: 1957–1973.e1956

Li CJ, Xiao Y, Yang M, Su T, Sun X, Guo Q, Huang Y, Luo XH (2018) Long noncoding RNA Bmncr regulates mesenchymal stem cell fate during skeletal aging. J Clin Invest 128: 5251–5266

Love MI, Huber W, Anders S (2014) Moderated estimation of fold change and dispersion for RNA-seq data with DESeq2. Genome Biol 15: 550

Lyabin DN, Eliseeva IA, Ovchinnikov LP (2014) YB-1 protein: functions and regulation. Wiley Interdiscip Rev RNA 5: 95–110

McCarthy I (2006) The physiology of bone blood flow: a review. J Bone Joint Surg Am 88 Suppl 3: 4–9

Mordovkina D, Lyabin DN, Smolin EA, Sogorina EM, Ovchinnikov LP, Eliseeva I (2020) Y-Box Binding Proteins in mRNP Assembly, Translation, and Stability Control. Biomolecules 10

Pan Y, Jiang Z, Ye Y, Zhu D, Li N, Yang G, Wang Y (2023) Role and Mechanism of BMP4 in Regenerative Medicine and Tissue Engineering. Ann Biomed Eng

Peng H, Hu B, Xie LQ, Su T, Li CJ, Liu Y, Yang M, Xiao Y, Feng X, Zhou R et al (2022) A mechanosensitive lipolytic factor in the bone marrow promotes osteogenesis and lymphopoiesis. Cell Metab 34: 1168–1182.e1166

Pittenger MF, Mackay AM, Beck SC, Jaiswal RK, Douglas R, Mosca JD, Moorman MA, Simonetti DW, Craig S, Marshak DR (1999) Multilineage potential of adult human mesenchymal stem cells. Science 284: 143–147

Rachner TD, Khosla S, Hofbauer LC (2011) Osteoporosis: now and the future. Lancet 377: 1276–1287

Rafii S, Butler JM, Ding BS (2016) Angiocrine functions of organ-specific endothelial cells. Nature 529: 316–325

Ramasamy SK, Kusumbe AP, Wang L, Adams RH (2014) Endothelial Notch activity promotes angiogenesis and osteogenesis in bone. Nature 507: 376–380

Sadat Tabatabaei Mirakabad F, Nejati-Koshki K, Akbarzadeh A, Yamchi MR, Milani M, Zarghami N, Zeighamian V, Rahimzadeh A, Alimohammadi S, Hanifehpour Y, et al (2014) PLGA-based nanoparticles as cancer drug delivery systems. Asian Pac J Cancer Prev 15: 517–535

Smith DM, Khairi MR, Johnston CC, Jr. (1975) The loss of bone mineral with aging and its relationship to risk of fracture. J Clin Invest 56: 311–318

Smith MR, Costa G (2022) RNA-binding proteins and translation control in angiogenesis. Febs j 289: 7788–7809

Suh KS, Lee YS, Kim YS, Choi EM (2013) Sciadopitysin protects osteoblast function via its antioxidant activity in MC3T3-E1 cells. Food Chem Toxicol 58: 220–227

Takahashi M, Shimajiri S, Izumi H, Hirano G, Kashiwagi E, Yasuniwa Y, Wu Y, Han B, Akiyama M, Nishizawa S et al (2010) Y-box binding protein-1 is a novel molecular target for tumor vessels. Cancer Sci 101: 1367–1373

Tikhonova AN, Dolgalev I, Hu H, Sivaraj KK, Hoxha E, Cuesta-Domínguez Á, Pinho S, Akhmetzyanova I, Gao J, Witkowski M et al (2019) The bone marrow microenvironment at single-cell resolution. Nature 569: 222–228

Tomlinson RE, Silva MJ (2013) Skeletal Blood Flow in Bone Repair and Maintenance. Bone Res 1: 311–322

Tuckermann J, Adams RH (2021) The endothelium-bone axis in development, homeostasis and bone and joint disease. Nat Rev Rheumatol 17: 608–620

Tunçay M, Caliş S, Kaş HS, Ercan MT, Peksoy I, Hincal AA (2000) Diclofenac sodium incorporated PLGA (50:50) microspheres: formulation considerations and in vitro/in vivo evaluation. Int J Pharm 195: 179–188

Wang L, Zhou F, Zhang P, Wang H, Qu Z, Jia P, Yao Z, Shen G, Li G, Zhao G et al (2017) Human type H vessels are a sensitive biomarker of bone mass. Cell Death Dis 8: e2760

Wu R, Feng S, Li F, Shu G, Wang L, Gao P, Zhu X, Zhu C, Wang S, Jiang Q (2023) Transcriptional and post-transcriptional control of autophagy and adipogenesis by YBX1. Cell Death Dis 14: 29

Xiao Y, Cai GP, Feng X, Li YJ, Guo WH, Guo Q, Huang Y, Su T, Li CJ, Luo XH et al (2023) Splicing factor YBX1 regulates bone marrow stromal cell fate during aging. Embo j: e111762

Xiao YZ, Yang M, Xiao Y, Guo Q, Huang Y, Li CJ, Cai D, Luo XH (2020) Reducing Hypothalamic Stem Cell Senescence Protects against Aging-Associated Physiological Decline. Cell Metab 31: 534–548.e535

Xie H, Cui Z, Wang L, Xia Z, Hu Y, Xian L, Li C, Xie L, Crane J, Wan M et al (2014) PDGF-BB secreted by preosteoclasts induces angiogenesis during coupling with osteogenesis. Nat Med 20: 1270–1278

Xu D, Xu S, Kyaw AMM, Lim YC, Chia SY, Chee Siang DT, Alvarez-Dominguez JR, Chen P, Leow MK, Sun L (2017) RNA Binding Protein Ybx2 Regulates RNA Stability During Cold-Induced Brown Fat Activation. Diabetes 66: 2987–3000

Xu R, Yallowitz A, Qin A, Wu Z, Shin DY, Kim JM, Debnath S, Ji G, Bostrom MP, Yang X et al (2018) Targeting skeletal endothelium to ameliorate bone loss. Nat Med 24: 823–833

Xue X, Huang J, Yu K, Chen X, He Y, Qi D, Wu Y (2020) YB-1 transferred by gastric cancer exosomes promotes angiogenesis via enhancing the expression of angiogenic factors in vascular endothelial cells. BMC Cancer 20: 996

Yang Y, Wang L, Han X, Yang WL, Zhang M, Ma HL, Sun BF, Li A, Xia J, Chen J et al (2019) RNA 5-Methylcytosine Facilitates the Maternal-to-Zygotic Transition by Preventing Maternal mRNA Decay. Mol Cell 75: 1188–1202.e1111

Zuo H, Yang D, Wan Y (2021) Fam20C Regulates Bone Resorption and Breast Cancer Bone Metastasis through Osteopontin and BMP4. Cancer Res 81: 5242–5254

